# Establishing MS2-MCP-based single-molecule RNA visualization in *Schizosaccharomyces pombe*

**DOI:** 10.64898/2026.03.09.710516

**Authors:** Douglas E. Weidemann, Sarah C. Turner, Samir G. Chethan, Silke Hauf

## Abstract

Single-molecule RNA imaging using the MS2–MCP system has transformed the study of RNA biology across model organisms. However, this technology has remained unavailable for fission yeast *(Schizosaccharomyces pombe)*, even though fission yeast is a central model for eukaryotic gene expression. Achieving single-molecule sensitivity requires identifying a narrow optimum where RNA labels are sufficiently bright while background fluorescence remains minimal. We have now accomplished this for *S. pombe* by systematically optimizing MCP expression and localization—screening a panel of constitutive *S. pombe* promoters and evaluating combinations of nuclear localization and export signals (NLSs and NESs). The resulting, successful constructs use tandem StayGold as the MCP fluorescent tag, taking advantage of its superior photostability. Together with optimized vectors for MS2 stem-loop tagging of endogenous transcripts, these tools enable single-molecule RNA imaging in fission yeast, opening the door to quantitative analyses of RNA dynamics in this core genetic model.

**Summary statement:** We establish single-molecule RNA visualization with MS2/MCP for fission yeast, *S. pombe*, an important model organism to study RNA biology.

## Introduction

The visualization of RNA molecules in live cells provides critical insights into their spatial and temporal dynamics (Gerber et al., 2023; Le et al., 2022; Tutucci et al., 2018). Such visualization has been achieved by repurposing RNA stem-loop-forming sequences from bacteriophages (Bertrand et al., 1998). The first, and still most widely used, version comes from the phage MS2. RNAs that carry MS2 stem-loops can be visualized with fluorescently tagged MS2 coat protein (MCP) (**Fig. 1A**). This system has enabled imaging of single RNA molecules with high spatial and temporal resolution and has, for example, led to key discoveries on the molecular basis of transcription bursts (Chubb et al., 2006; Donovan et al., 2019; Fukaya et al., 2016; Golding et al., 2005; Patel et al., 2023), splicing (Brody et al., 2011; Coulon et al., 2014; Martin et al., 2013; Schmidt et al., 2011), RNA nuclear export dynamics (Grünwald and Singer, 2010; Mor et al., 2010), RNA localization (Bertrand et al., 1998; Fusco et al., 2003), translation (Halstead et al., 2015; Morisaki et al., 2016; Pichon et al., 2016; Wu et al., 2016) and RNA degradation (Horvathova et al., 2017).

**Figure 1.**
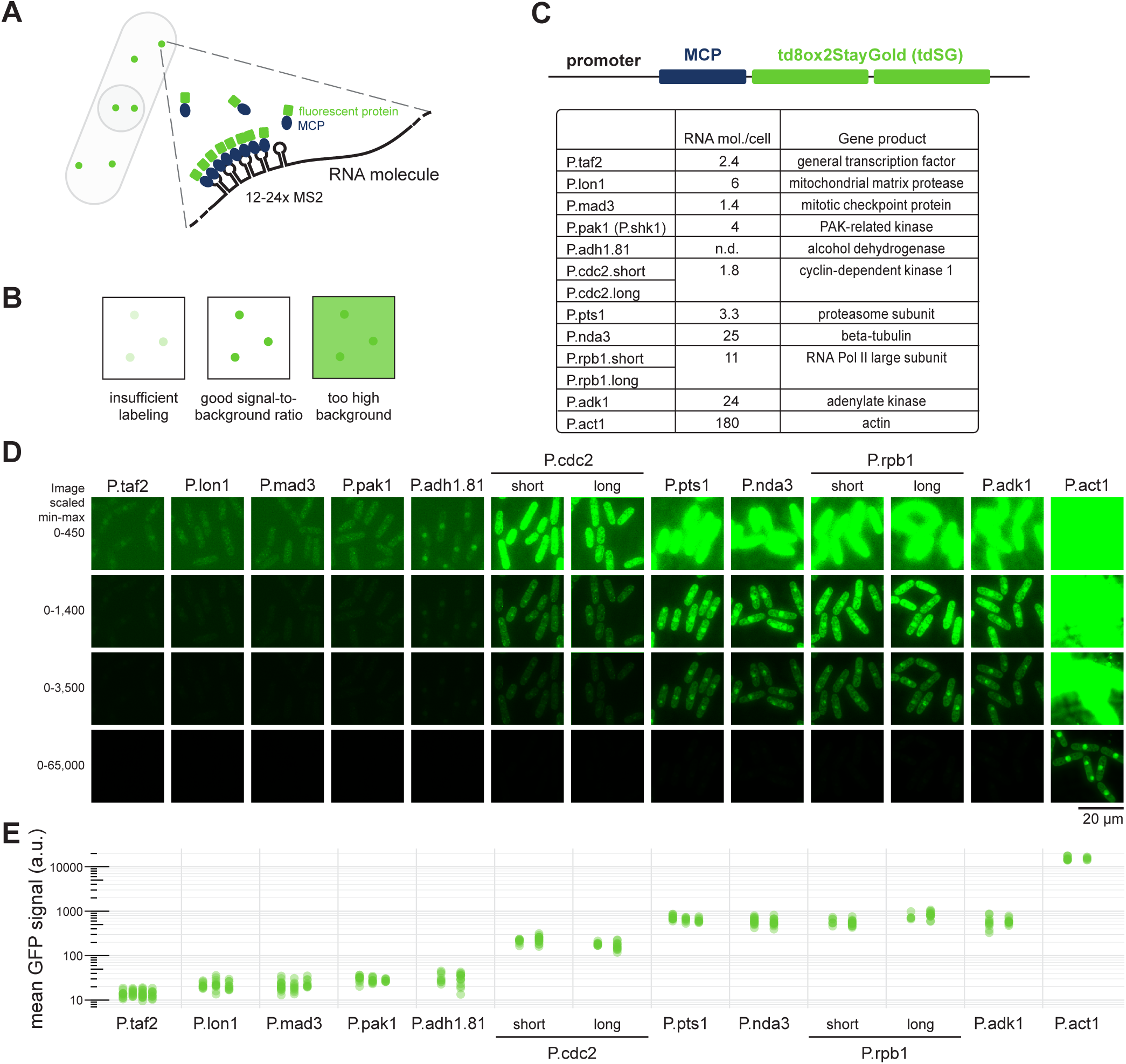
Screening constitutive *S. pombe* promoters to establish RNA imaging with MS2-MCP. **(A)** RNAs are tagged with stem-loops from the bacteriophage MS2. Dimers of MS2 coat protein (MCP) bind to each stem-loop structure. Tagging of MCP with a fluorescent protein allows for visualization of the RNA. **(B)** RNA single-molecule imaging requires a high enough MCP-fluorescent protein concentration to efficiently label the RNAs, but low enough concentration to not create a high background signal. **(C)** MCP was tagged with tandem StayGold (tdSG), a highly photobleaching-resistant green fluorescent protein and expressed under different promoters from the *S. pombe ura4* locus. To optimize the signal-to-background ratio, 11 *S. pombe* promoters were tested. The estimates of RNA molecules per cell for each respective gene are from Marguerat *et al*., Cell 2012. For P.cdc2 and P.rpb1 both a shorter and a longer sequence upstream of the gene’s start codon was tested. **(D)** Example images from strains expressing MCP-NLS-tdSG from the indicated promoters. Cells also express *mad2*-24xMS2. Image acquisition conditions were the same; each field of view is shown using four different scaling settings in order to capture the breadth of signal intensities. **(E)** The mean tdSG signal intensity in single cells was quantified from at least two different images for each strain (between 5 and 16 cells per image); dots are individual cells. Note that data are displayed on a logarithmic scale.

Fission yeast (*Schizosaccharomyces pombe*) is a widely used model organism in molecular biology, including RNA biology (Hoffman et al., 2015; Vyas et al., 2021). Research using *S. pombe* has uncovered fundamental mechanisms of transcription, splicing, nuclear and cytoplasmic RNA decay, and RNA-mediated mechanisms of heterochromatin formation (Fair and Pleiss, 2017; Hirota et al., 2008; Lee et al., 2013; Schramke et al., 2005; Sugiyama and Sugioka-Sugiyama, 2011; Ukleja et al., 2016; Vo et al., 2019; Volpe et al., 2002; Yan et al., 2015). However, RNA imaging with single-molecule sensitivity in living cells has not yet been achieved in *S. pombe*. A few *S. pombe* RNAs have been labelled fluorescently using MS2-MCP, but only to visualize bulk nuclear versus cytoplasmic localizations (Carnahan et al., 2005) or an abundantly transcribed RNA at its transcription site (Shichino et al., 2014).

Critical for single-molecule sensitivity is the expression level and localization of MCP. Expression that is too weak does not yield enough signal from each tagged RNA molecule, whereas overly high expression floods the cell with unbound fluorescent MCP, impairing the visualization of labelled RNA molecules against the high background (**Fig. 1B**). Screening a broad range of constitutive *S. pombe* promoters and several localization signal combinations, we have now identified MCP constructs suitable for single-molecule RNA visualization in *S. pombe*. These constructs use the highly photobleaching-resistant green fluorescent protein StayGold fused to MCP (Ando et al., 2023; Hirano et al., 2022), thus making use of the latest advancements in fluorescent protein technology.

## Results

### Identification of constitutive *S. pombe* promoters suitable for single-molecule MS2-MCP imaging

To optimize the expression of MCP, we selected constitutive *S. pombe* promoters based on gene function (*pts1, nda3*), known low-noise expression (*mad3*, *rpb1* (Weidemann et al., 2023)), stable expression across stress conditions (*cdc2, taf2, lon1, adk1* (Thodberg et al., 2019)), or their prior use in expression vectors (*adh1.81, pak1, act1* (Chen et al., 2017; Kiriya et al., 2017; Vještica et al., 2019; Yokobayashi and Watanabe, 2005)) (**Fig. 1C**). These promoters span a range of native mRNA expression levels, from around 1 to 180 mRNA molecules per cell (Marguerat et al., 2012). As promoter region, we used between 214 and 913 base pairs upstream of the respective start codon, encompassing both the promoter and the 5’UTR of each gene. Using these promoter regions, we expressed MCP fused to tandem StayGold (td8ox2StayGold, tdSG (Ando et al., 2023)) (**Fig. 1D,E**). StayGold was chosen due to its superior photostability compared to other fluorescent proteins (Ando et al., 2023; Hirano et al., 2022). We observed four classes of fluorescence intensities, partly, but not perfectly, scaling with the reported RNA numbers: low *(taf2, lon1, mad3, pak1, adh1.81)*, medium *(cdc2)*, high *(pts1, nda3, rpb1, adk1),* and very high expression *(act1)* (**Fig. 1E**, note the logarithmic scale). Monomeric StayGold (Ando et al., 2023), tested as an alternative to tdSG, yielded signals much weaker than half the tdSG signal, was undetectable with weak promoters, and therefore was not pursued further (**Fig. S1**).

To test the feasibility of imaging single RNA molecules, we replaced the wild-type *mad2* gene (a component of the spindle assembly checkpoint (McAinsh and Kops, 2023)) with a version containing a C-terminal EGFP tag that was made non-fluorescent by the Y66L mutation (Rosenow et al., 2004) and 24 MS2 stem-loops in the 3’UTR (*mad2*-24xMS2). By single-molecule RNA fluorescence *in situ* hybridization (smFISH), the *mad2* gene shows between 0 and 7 mRNA molecules per cell (mean ∼ 2–3) (Esposito et al., 2022; Weidemann et al., 2023). We used the MS2V6 variant for tagging, which has slightly weakened MCP-binding and a longer linker between stem-loops to avoid an artificial stabilization of the MS2 fragment (Tutucci et al., 2017). To facilitate endogenous tagging, we slightly modified existing 12x MS2V6 and 24x MS2V6 vectors (**Fig. 2A**). The original vectors were designed as PCR templates to attach homology regions to the MS2 repeats; however, PCR across repeat regions often requires optimization (Hommelsheim et al., 2014). We therefore opted for a strategy where homology regions can be cloned into the vector upstream and downstream of the repeats, and the piece to be transformed into yeast is excised using Type IIS restriction enzymes (**Fig. 2, S2**). The donor DNA containing MS2 tags can be introduced into the genome in a scarless manner by CRISPR/Cas9-mediated homologous recombination (**Fig. 2B**) or by replacement of a counterselectable cassette (**Fig. 2C**). Alternatively, the kanamycin(G418)-resistance on the vector can be used (**Fig. 2D**).

**Figure 2.**
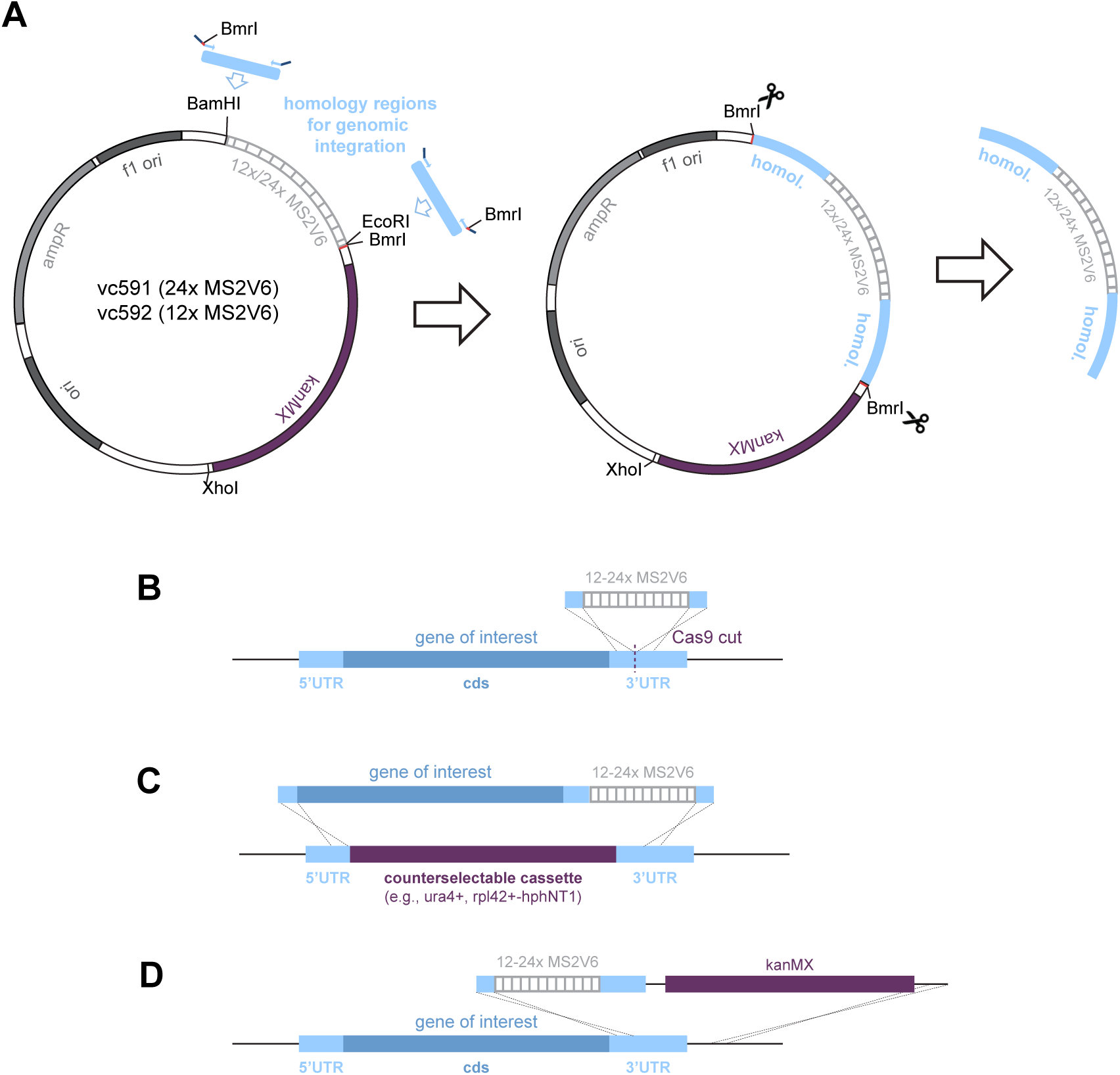
Vectors for attaching homology regions to MS2V6 repeats. **(A)** Vectors to append homology regions upstream and downstream of the MS2V6 repeats; vc591 (24x MS2V6) and vc592 (12x MS2V6) are slightly modified versions of Addgene plasmids # 104393 and # 104392 (Tutucci et al., Nat Methods 2017). An EcoRI site was added downstream of the MS2V6 repeats. Homology regions can be integrated at the BamHI, EcoRI, and XhoI sites. **(B,C,D)** Strategies to integrate the 12-24x MS2V6 repeats into the genome: (B) CRISPR/Cas9-facilitated integration without resistance gene; (C) replacement of a counterselectable cassette, e.g., when working with a non-essential gene; (D) integration using the kanMX resistance gene present on the vector.

Using the combination of *mad2-24xMS2* with MCP-tdSG expressed from different promoters, we observed clear dot-like signals in the cytoplasm using the *lon1*, *mad3*, *pak1*, and *cdc2* promoters (**Fig. 3A,B, S3**). The abundance and movement of these dots are consistent with the expected behavior of individual mRNA molecules (**Fig. 3B, S3, Supplemental Movies S1-S4**). Such signals were not observed when MCP-tdSG was expressed in the absence of an MS2 tag (**Fig. 3B, Supplemental Movies S1-S4**). As expected, promoters that were too weak *(taf2)* or too strong (*pts1* or stronger) failed to produce clearly discernible RNA signals (**Fig. S3**). Similar ratios of RNA signals over cytoplasmic background were obtained with MCP-tdSG expressed from the *lon1*, *mad3*, *pak1*, and *cdc2* promoters, despite the stronger MCP-tdSG expression from the *cdc2* promoter (**Fig. 3C, S4)**, indicating that the MS2 stem-loops are not saturated with MCP when using weaker promoters.

**Figure 3.**
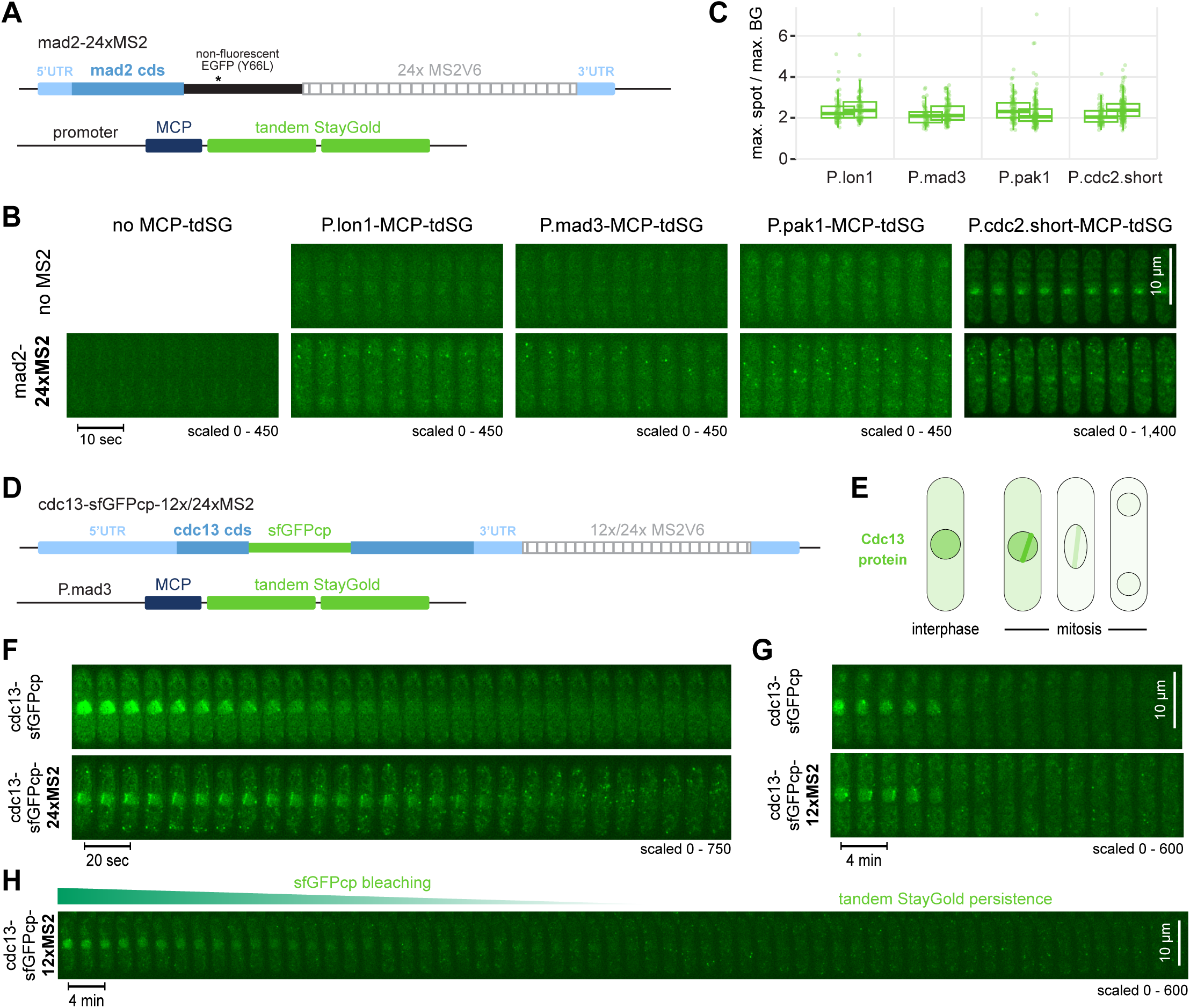
Visualization of single RNA molecules with MCP-tdSG expressed from different promoters. **(A)** The *mad2* gene was tagged C-terminally with a non-fluorescent version of EGFP and the 24x MS2V6 cassette was inserted in the 3’UTR. MCP-tdSG was expressed from different promoters. **(B)** Short kymographs from live-cell imaging; longer sequences are shown in Fig. S3. For each MCP-tdSG construct, a strain without integration of MS2 repeats is shown as control. Far left: strain expressing *mad2*-24xMS2, but no MCP-tdSG. Images are maximum intensity projections of the Z-stack. Note the different scaling setting for P.cdc2.short-MCP-tdSG. **(C)** Quantification of the maximum spot intensity over maximum background intensity. Two replicates per strain, 56-162 spots per replicate. Points: individual spots; box plot: summary statistics (center line, median; box boundaries, 1st and 3rd quartiles). **(D)** The *cdc13* gene was internally tagged with circularly permuted superfolder GFP (sfGFPcp) and a 12x or 24x MS2V6 cassette was inserted in the 3’UTR. The construct was expressed under endogenous *cdc13* regulatory sequences (promoter, terminator) from the *leu1* locus. MCP-tdSG was expressed from the *mad3* promoter. **(E)** Schematic illustrating Cdc13 protein localization. Cdc13 accumulates during interphase and is strongly enriched in the nucleus. Cdc13 localizes to spindle pole bodies and the spindle during early mitosis and becomes degraded at the metaphase-to-anaphase transition. **(F,G)** Kymographs from live-cell imaging of cells undergoing mitosis; *cdc13*-sfGFP tagged with either 24x MS2V6 (F) or 12x MS2V6 (G). A strain without MS2 repeats is shown as control. Images were recorded every 5 sec (F) or 12 sec (G); every second image (10 sec, F) or every 10th image (2 min, G) is shown. Images are maximum intensity projections of the Z-stack. **(H)** Similar to (G), but showing an interphase cell; loss of the nuclear sfGFPcp signal is due to photobleaching, not degradation.

To test a second gene with higher mRNA concentration, we tagged the cyclin *cdc13* with MS2 stem-loops (**Fig. 3D**). Cdc13 mRNA numbers per cell determined by smFISH range from around 10 to 35 (mean ∼ 20) (Bashir et al., 2026; Vandal et al., 2026; Weidemann et al., 2023). In addition to the MS2 tag in the 3’UTR, the *cdc13* gene was internally tagged with circularly permuted, superfolder GFP (sfGFPcp), so that Cdc13-sfGFPcp protein could be visualized in addition to *cdc13* mRNA. Because the Cdc13 protein enriches strongly in the nucleus (Alfa et al., 1989; Decottignies et al., 2001), the Cdc13-sfGFPcp protein signal is expected to only contribute minimally to background in the cytoplasm (**Fig. 3E**). With MCP-tdSG expressed from the *mad3* promoter, both Cdc13 protein, which is degraded in mitosis, and *cdc13* mRNA could be tracked (**Fig. 3F,G, S5**). No obvious difference in Cdc13-sfGFPcp protein signals was observed with or without the MS2 tags, suggesting that the mRNA with MS2 stem-loops remained functional. Dot-like signals in the cytoplasm, consistent with single mRNA molecules, were observed both with a 12x and 24xMS2 tag, but not when *cdc13* was expressed without MS2 tag (**Fig. 3F,G, S5, Supplemental Movies S5,S6**). MCP-tdSG dot-like signals in the cytoplasm could still be monitored once the Cdc13-sfGFPcp nuclear signal had bleached (**Fig. 3H**), consistent with the expected photobleaching-resistance of tdSG (Ando et al., 2023; Hirano et al., 2022).

Taken together, we have identified several constitutive promoters that allow for single-molecule mRNA imaging in *S. pombe* using MS2 tags and MCP-tdSG. The signals are sufficiently prominent to be observable even with some additional green fluorescent protein background, and tdSG confers considerable bleaching resistance.

### MS2-tagged mRNAs are mostly intact, and numbers are in the wild-type range

In some instances, MS2 tags have been observed to impair the degradation of the tagged RNA or to remain undegraded after degradation of the rest of the RNA molecule (Garcia and Parker, 2015; Garcia and Parker, 2016; Haimovich et al., 2016; Heinrich et al., 2016). To assess whether this is a concern, we turned to single-molecule mRNA FISH in fixed cells and labelled both the MS2 repeats and the coding sequence of the tagged RNA. For *mad2*-24xMS2, we performed double-labelling with CAL610-tagged probes against the non-fluorescent EGFP tag and Quasar570-tagged probes against MS2V6 (**Fig. 4A,B, S6A-C**). We found that the probes against the coding sequence and the MS2V6 tag largely co-localized (**Fig. 4B,C**). Perfect co-localization is not expected because of the exonuclease-mediated digestion of mRNA, which in the yeast cytoplasm occurs predominantly in 5’ to 3’ direction (Grochowski et al., 2024; Parker, 2012). The mRNA numbers per cell detected with either the EGFP or the MS2 probes (mean ∼2.3, **Fig. 4D**) were consistent with previous measurements for EGFP-tagged and untagged *mad2* (mean 2.3 and 2.7 (Esposito et al., 2022)). For *cdc13*, we performed separate single-labelling with Quasar570-tagged probes against the sfGFPcp coding sequence or against MS2V6 (**Fig. 4E,F**). We again found similar mRNA numbers per cell obtained with both probes (mean 27.5 and 26.2, **Fig. 4G**). Furthermore, the numbers were in the same range as the numbers for *cdc13*-sfGFPcp mRNA without MS2 tag (mean 27.5, **Fig. S6D**) (Bashir et al., 2026; Vandal et al., 2026; Weidemann et al., 2023).

**Figure 4.**
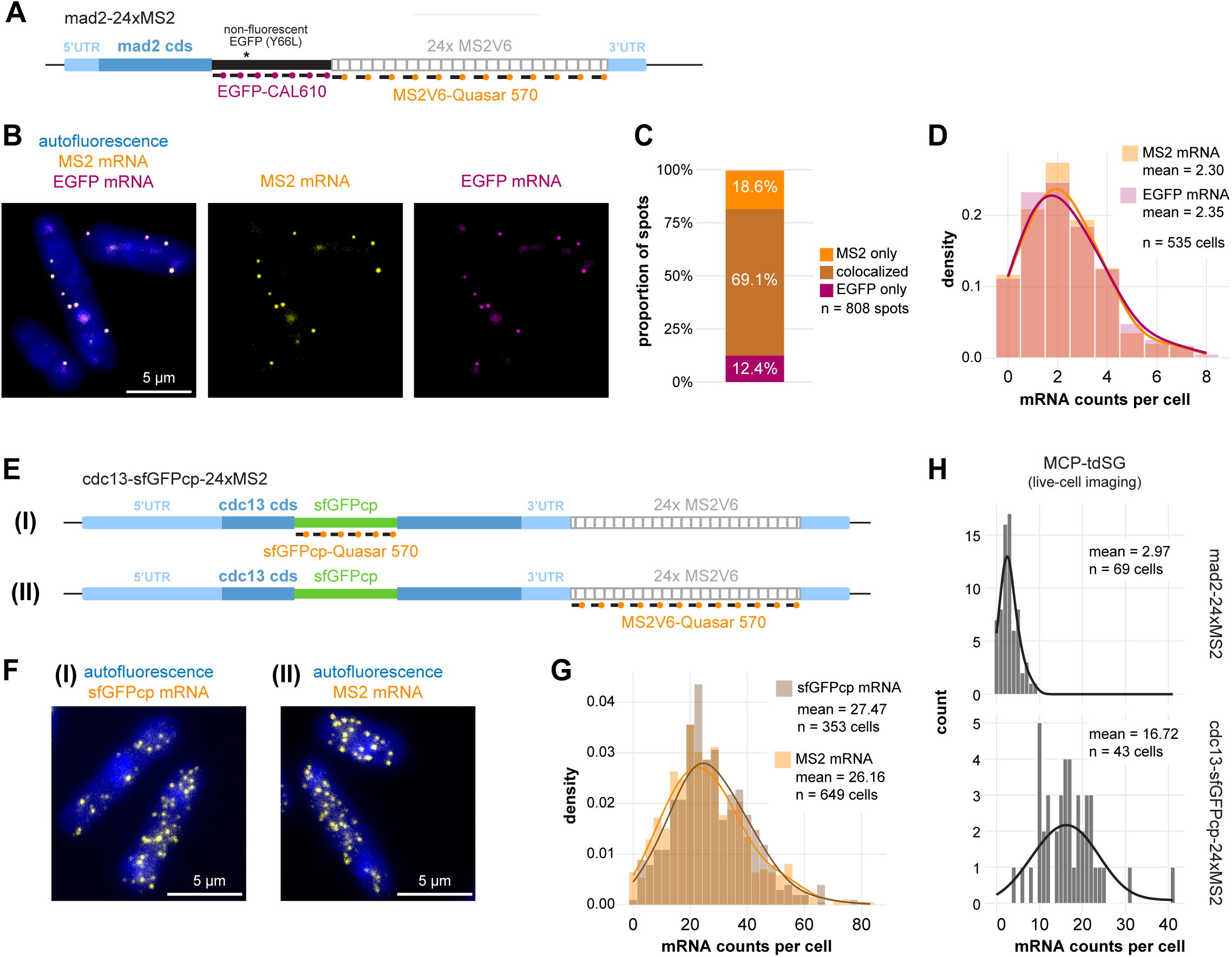
MS2 probes largely co-localize with probes against the coding sequence, and both yield similar mRNA numbers. **(A)** Schematic of two-color RNA FISH with probes targeting the EGFP tag and the 24x MS2 tag in *mad2*-darkEGFP-24xMS2 cells. **(B)** Representative images (maximum intensity projection) of the experiment outlined in (A). **(C)** Proportion of spots co-localizing or not in the same experiment. **(D)** Counts of mRNA molecules per cell from the same experiment. Lines are Gaussian kernel density estimates (smoothing bandwidth x 2). **(E)** Schematic of single-color RNA FISH with probes targeting either the sfGFPcp tag or the 24x MS2 tag in *cdc13*-sfGFPcp-24xMS2 cells. **(F)** Representative images (maximum intensity projections) of the experiments outlined in (E). **(G)** Counts of mRNA molecules per cell from the experiments outlined in (E). Lines as in (D). **(H)** Counts of mRNA molecules per cell from RNA live-cell imaging, using the same strains as in the RNA FISH experiments. Lines as in (D).

Finally, we compared the numbers obtained by single-molecule mRNA FISH with those observed in mRNA live-cell imaging using MCP-tdSG. We found similar numbers in mRNA live-cell imaging as in the FISH experiments (**Fig. 4H**). The mean mRNA number per cell for *mad2*-24xMS2 was 2.97, that for *cdc13*-sfGFPcp-24xMS2 was 16.7. We suspect that the MCP-tdSG concentration may be slightly suboptimal for the larger number of *cdc13* mRNA molecules and that live-cell imaging for *cdc13* could be improved by raising the MCP-tdSG concentration.

Taken together, these results indicate that the MS2-tagged mRNAs retain the expression profile of the corresponding untagged mRNA, that mRNA degradation is not greatly impaired by the MS2V6 tag, and that detection by MCP-tdSG is efficient.

### Cell cycle-dependent mRNA expression can be visualized in live cells

To further validate that mRNA dynamics are not strongly impaired by the MS2 tag, we tagged *rad21*, a gene that codes for one of the subunits of the cohesin complex (**Fig. 5A**) (Tomonaga et al., 2000). The *rad21* gene is expressed in a cell cycle-dependent manner (Marguerat et al., 2006; Oliva et al., 2005; Peng et al., 2005; Rustici et al., 2004). In *S. pombe*, *rad21* mRNA levels are highest just prior to cell division (Weidemann et al., 2023). Live-cell imaging of *rad21-*24xMS2 cells reproduced this pattern and showed a peak of mRNA during an approx. 30 min period just prior to septation and cell division (**Fig. 5B,C; Supplemental Movie S7**). The sharp increase and decrease of mRNA numbers corroborate that transcription and mRNA degradation are likely not majorly affected by the presence of the MS2V6 stem-loops.

**Figure 5.**
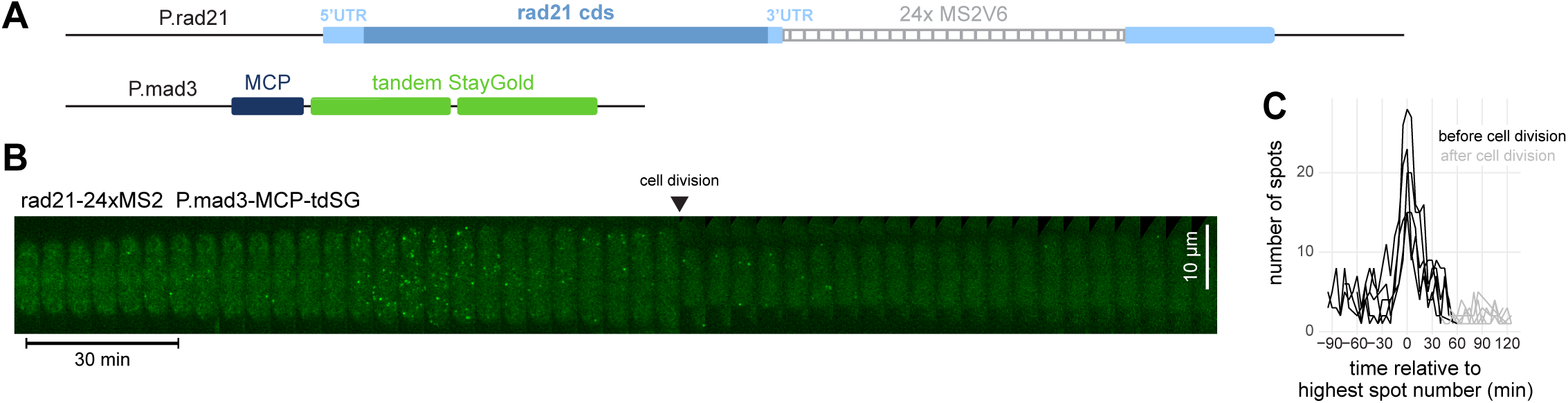
Visualization of the cell cycle-dependent expression of cohesin (*rad21*-24xMS2). **(A)** The *rad21* gene was tagged in the 3’UTR with 24x MS2V6 and was integrated into the genome at the *ade6* locus using its own promoter (P.rad21). MCP-tdSG was expressed from the *ura4* locus using the *mad3* promoter. **(B)** Representative kymograph from live-cell imaging. A Z-stack was recorded every 5 min; shown are maximum intensity projections of the Z-stack. **(C)** Quantification of spot numbers in 5 cells using U-FISH. Curves were aligned to the maximum number measured. Cell division is indicated by the transition from black to gray line.

### Visualization of cytoplasmic mRNAs benefits from a weak NLS or a combination of NLS and NES

MCP-fluorescent protein fusion constructs often include nuclear localization sequences (NLSs) to minimize background of unbound MCP in the cytoplasm, although the argument has also been made that omitting an NLS may be beneficial (Tocchini and Mango, 2024). Our initial MCP-tdSG version included an SV40 NLS with minimal surrounding sequences (**Fig. 6A**). Using this construct, MCP was visible in the cytoplasm, and the level of nuclear enrichment varied with the promoter and 5’UTR used (**Fig. S3**). With many of the promoter and 5’UTR combinations, MCP-tdSG was further enriched in the nucleolus (**Fig. S3**). Expression of a tandem MCP construct (stdMCP (Wu et al., 2015)) with the same SV40 NLS, but a longer linker upstream of the NLS (stdMCP-NLS*-tdSG) yielded a much stronger nuclear enrichment—so strong that it became inefficient in labelling cytoplasmic mRNAs (**Fig. 6, S7A**). Shortening the linker between MCP and NLS increased the fraction of stdMCP-tdSG in the cytoplasm again (**Fig. S7A**), indicating that the efficiency of the SV40 NLS is compromised by its close vicinity to MCP, but that this may be beneficial for cytoplasmic mRNA imaging. For a better overview on the influence of nucleocytoplasmic distribution on RNA imaging, we additionally tested tandem NLSs with either one or two PKI nuclear export signals (NESs) (**Fig. 6A**). Addition of a second NLS strongly increased the nuclear enrichment, similar to the single NLS with longer linker, and also decreased the intensity of RNA signals in the cytoplasm (**Fig. 6B**). Addition of one or two NESs progressively increased the cytoplasmic and decreased the nuclear signal, bringing back cytoplasmic RNA labelling (**Fig. 6B**). Thus, the nucleocytoplasmic ratio is well tuneable by different NLS and NES combinations. Expression of these constructs from two different promoters *(P.cdc2* and *P.mad3)* yielded qualitatively similar results (**Fig. S7C,D**).

**Figure 6.**
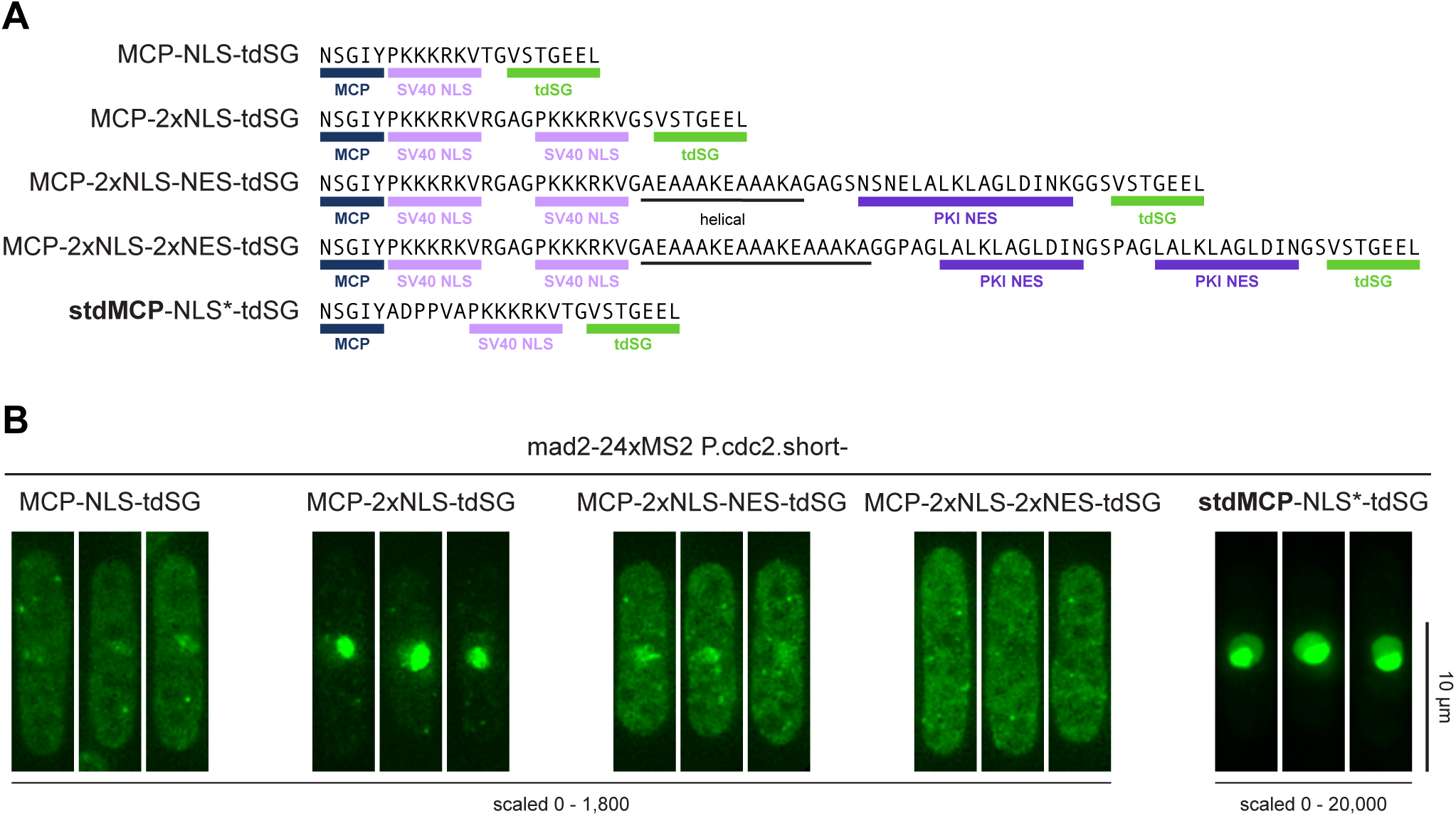
Spatial distribution of MCP-tdSG can be tuned by different combinations of NLS and NES sequences. **(A)** Overview of NLS and NES combinations. NLS and NLS* differ by the linker sequence between MCP and the SV40 NLS. **(B)** Example images from strains expressing *mad2*-24xMS2 and the indicated MCP-tdSG or stdMCP-tdSG constructs. MCP constructs were expressed from the *cdc2* promoter. See Fig. S7 for the same constructs expressed from the *mad3* promoter. All images were recorded with the same exposure conditions; maximum intensity projections of Z-stacks are shown.

In summary, we have identified conditions for single-molecule mRNA imaging in *S. pombe*, which, for the constructs tested, seemed to preserve mRNA functionality. We provide a range of integration vectors for expression of MCP-tdSG that can be paired with MS2-tagged genes (**Fig. 7**). These vectors use the pUra4AfeI backbone (Vještica et al., 2019) for stable integration into the *S. pombe ura4* locus. These tools now set the stage for single-molecule-based exploration of RNA biology using *S. pombe*.

**Figure 7.**
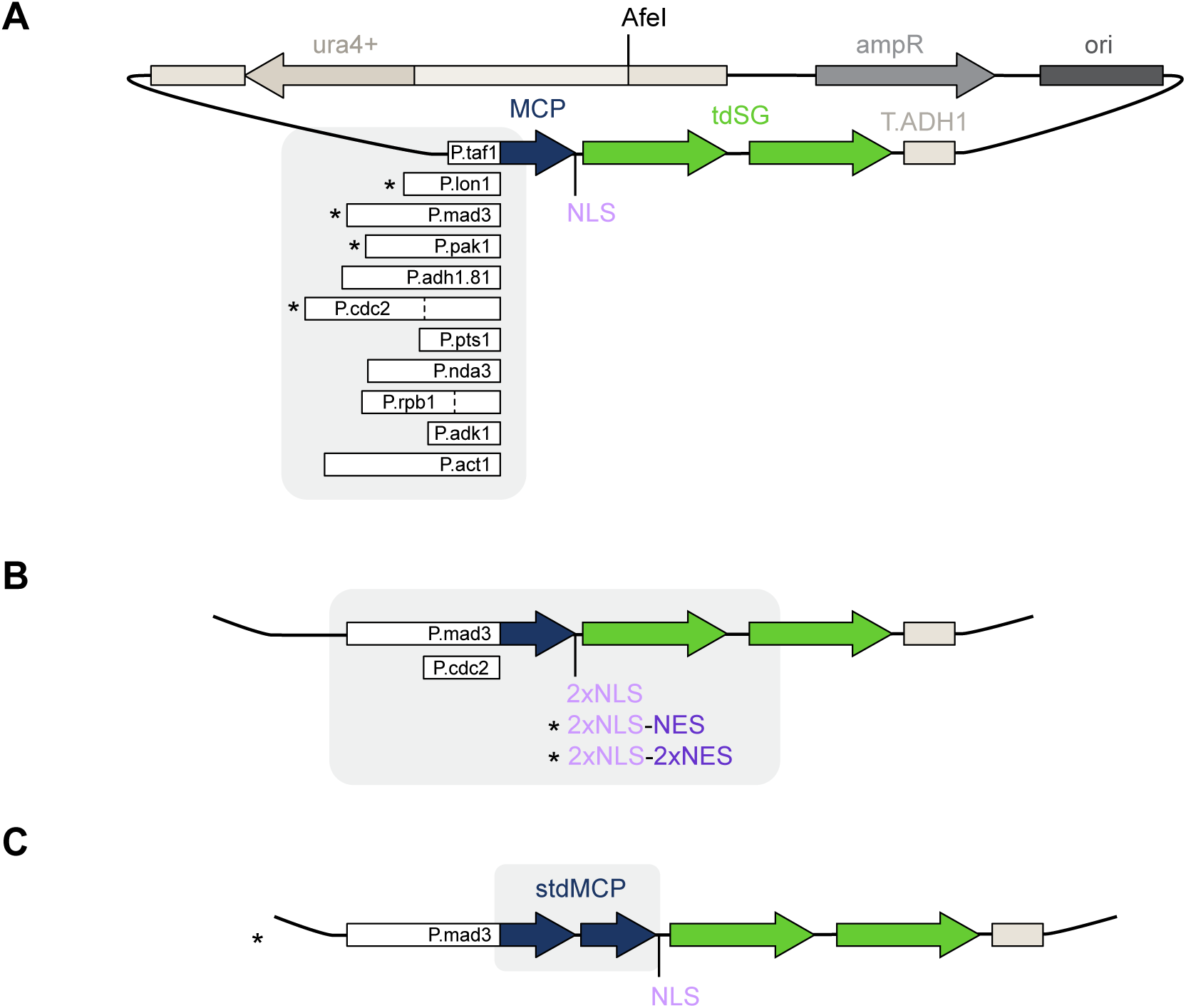
MCP-tdSG vectors for *S. pombe* expression. All vectors are derivatives of pUra4AfeI (Vjestica et al., 2019) and can be integrated at the ura4 locus after linearization of the vector with AfeI. Vectors marked with an asterisk have been deposited at Addgene and RIKEN. Other vectors are available upon request. **(A)** Vectors for expression of MCP-NLS-tdSG from different constitutive promoters. The dashed lines for P.cdc2 and P.rpb1 indicate the length of the short promoter versions. **(B)** Vectors for expression of MCP-tdSG with different NLS/NES combinations from either the *mad3* or the short *cdc2* promoter. **(C)** Vector for expression of synonymized tandem MCP (stdMCP, Wu et al., 2015), tagged with NLS and tdSG.

## Discussion

The MS2-MCP system is widely used for single-molecule RNA live-cell imaging. The absence of its implementation for the model organism *S. pombe* was a notable gap. Here, we close this gap by establishing *S. pombe* expression vectors for MCP fused to bleaching-resistant tandem StayGold that yield cytoplasmic concentrations suitable for MS2 imaging (**Fig. 7**). By combining MCP-tdSG with several NLS/NES combinations, we provide further flexibility in tuning the nuclear and cytoplasmic signals (**Fig. 6,7**). We also modified available MS2 vectors (Tutucci et al., 2017) to facilitate MS2 tagging (**Fig. 2**). Collectively, these tools now enable the exploration of RNA life cycle dynamics and RNA localization with single-molecule precision in *S. pombe* cells. Our data also provide insights into the expression strengths achievable with different constitutive promoters (**Fig. 1**), which may be informative for other applications; however, we caution that expression strength is further modulated by the coding sequence. For example, in our experiments, the tandem MCP (stdMCP) construct, containing a codon-optimized sequence (Wu et al., 2015), yielded higher concentrations from the same promoter than the single-copy MCP (**Fig. S7**).

While the constructs we provide here serve as an excellent starting point, further adjustments are possible. On the MCP side, tandem MCP has been reported to provide more consistent signals than single-copy MCP, presumably because MCP acts as a dimer and the tandem construct is poised for dimerization (Wu et al., 2012; Wu et al., 2015). We observed that expression of tandem stdMCP-tdSG resulted in an overall stronger signal and stronger nuclear enrichment than tdMCP-tdSG (**Fig. S7**). Its usefulness can likely be improved by employing a weaker promoter and introducing an NES sequence. Furthermore, it may be possible to increase the signal-over-background ratio by using versions of MCP that are unstable unless bound to an MS2 stem-loop (Kuffner et al., 2026).

On the MS2 side, it is important to note that repetitive stem-loop tags can alter RNA physiology (Garcia and Parker, 2015; Garcia and Parker, 2016; Haimovich et al., 2016; Heinrich et al., 2016; Li et al., 2022; Tocchini and Mango, 2024; Tutucci et al., 2017). Thus, depending on the application, MS2 tags may require further optimization. For instance, MS2-tagging can lead to RNA mislocalization (Tocchini and Mango, 2024), and MS2 stem-loops may persist in cells after the remainder of the tagged mRNA has been degraded (Garcia and Parker, 2015). These known challenges have prompted the optimization of MS2 tags for linker length, nucleotide composition, and MCP-binding strength (Katz et al., 2021; Tocchini and Mango, 2024; Tutucci et al., 2017; Wu et al., 2015). Here, we have used one of these optimized sequences, MS2V6 (Tutucci et al., 2017), but other optimized sequences may offer further benefits. For example, a version without start and stop codons will be beneficial for 5’ tagging (Halstead et al., 2015; Hocine et al., 2013), and versions with shortened linkers may minimize the targeting of tagged mRNAs by nonsense-mediated decay (NMD) (Tocchini and Mango, 2024).

The MS2-MCP system is not the only bacteriophage-derived system used for RNA tagging. The orthogonal PP7-PCP system is similarly popular (Gerber et al., 2023; Le et al., 2022; Tutucci et al., 2018). The insights gained here from testing different promoters and NLS/NES combinations will be transferable to this system, which further widens the opportunities to monitor and manipulate RNAs in *S. pombe*.

## Materials and Methods

### *S. pomb*e strains

All *S. pombe* strains are listed in Table S1. The *mad2* gene was tagged with a non-fluorescent (Y66L) version of EGFP (Rosenow et al., 2004) at its C-terminus and with 24x MS2V6 (Tutucci et al., 2017) immediately after the stop codon. The tagged *mad2* gene was integrated at the endogenous locus by first deleting the *mad2* coding sequence with a counterselectable *rpl42+/hygR* cassette in an *rpl42*-sP56Q background (Roguev et al., 2007) and then replacing the counterselectable cassette with *mad2-*darkEGFP-24xMS2V6 by selecting first for cycloheximide resistance and then confirming hygromycin sensitivity. The *cdc13* gene was tagged internally between amino acids S177 and V178 with circularly permuted superfolder GFP (sfGFPcp) (Kamenz et al., 2015) and after nucleotide 205 or 332 of the 3’UTR with 12x MS2V6 or 24x MSV6. The constructs were expressed from a pDUAL vector (Matsuyama et al., 2004) integrated at the exogenous *leu1* locus. No differences in functionality were observed between the integration sites in the 3’UTR. To construct the pDUAL vector, homology regions were first cloned into the 12x MS2V6 and 24x MS2V6 vectors, and a BamHI/XbaI fragment containing the partial coding sequence for *cdc13-*sfGFPcp and the 3’UTR with MS2 repeats was then cloned into a BamHI/SpeI-digested pDUAL vector containing cdc13-sfGFPcp expressed from the *cdc13* promoter. The pDUAL vector was linearized with NotI and integrated into the genome at the *leu1* locus by transformation into a *leu1-32* strain. The *rad21* gene was tagged with 24x MS2V6 70 nucleotides after the stop codon, and the entire gene, including 1,452 nucleotides of promoter sequence and 5’UTR, was integrated into a pAde6PmeI vector (Vještica et al., 2019). The vector was linearized with PmeI and integrated into the genome at the *ade6* locus by transformation into an *ade6-D16* strain. Integration of the vector restores *ade6+*. MCP-tdSG constructs were integrated by linearizing the corresponding pUra4AfeI vector with AfeI and transforming it into a *ura4+*-deleted strain (*ura4-D18*). Integration of the vector restores *ura4+*.

### Vectors

All vectors are listed in Table S2. Promoter sequences were amplified from the *S. pombe* genome, except for P.adh1.81, which was from the pRAD81 plasmid (gift from Yoshinori Watanabe). MCP-NLS was from pET296-YcpLac111 CYC1p-MCP-NLS-2xyeGFP (Addgene plasmid # 104394; gift from Robert Singer and Evelina Tutucci); stdMCP-NLS* was from pUbC-nls-ha-stdMCP-stdGFPx (Addgene plasmid # 98916; gift from Robert Singer); td8ox2StayGold was from pBS Coupler1/td8ox2StayGold (RIKEN RDB20227) and mStayGold from pRSETB/mStayGold (RIKEN RDB20214), both by Miyawaki et al. (Ando et al., 2023) and provided by the RIKEN BRC through the National BioResource Project of the MEXT, Japan. The ADH1 terminator from *S. cerevisiae* (*Scer\T.ADH1*) is the same sequence used in the *S. pombe* pFA6a expression vectors (Bähler et al., 1998). Additional NLS and NES sequences were inserted by digest and Gibson assembly with synthetic fragments.

The vectors containing 12x MSV6 (pET251-pUC 12xMS2V6 Loxp KANr Loxp, Addgene plasmid # 104392) and 24x MS2V6 cassettes (pET264-pUC 24xMS2V6 Loxp KANr Loxp, Addgene plasmid # 104393) were a gift from Robert Singer and Evelina Tutucci (Tutucci et al., 2017). They were modified to incorporate an EcoRI site downstream of the MS2 cassette by BglII digest and Gibson assembly with a synthetic fragment.

### Live-cell microscopy

Cells were grown in Edinburgh Minimal Medium (EMM, MP Biochemicals LLC, 114110012) with additional supplements (200 µg/mL leucine or 50 µg/mL uracil), as needed. For imaging, cells were diluted to 1 x 10^6 cells/mL in pre-warmed medium and 300 µL were mounted in one well of a glass-bottom 8-well µ-Slide (Ibidi, 80827) that had been coated with 50 µg/mL lectin (Sigma, L1395). For coating, 300 µL of lectin solution were added to wells for several hours at 30 °C; the solution was removed just prior to imaging, and the slide was air-dried at 30 °C. Imaging was performed on a DeltaVision widefield microscope, using an Olympus 60×/1.42 Plan APO oil objective, 461-489 nm LED illumination, a 525/48 nm GFP/FITC emission filter, a PCO edge sCMOS camera, and an environmental chamber to keep the temperature at 30 °C. Z-stacks were recorded every 5, 6, 12, or 300 sec, as indicated, and Z-sections were spaced by 0.3 to 0.5 µm over a distance of 3.9 to 5.4 µm. Imaging conditions were kept the same for samples and corresponding controls. Using these conditions, autofluorescence bleached in the first few frames, and those frames were removed. Images were deconvolved using SoftWoRx software with three cycles of the ratio method (conservative), noise filtering set to high (300 nm), a camera intensity offset of 0, and generally without corrections – except for the > 4h-hour imaging of *rad21*-24xMS2, where a bleaching correction was applied.

### Whole-cell fluorescence quantification

Cells were segmented based on the brightfield image using YeaZ (Dietler et al., 2020). Quantification was performed on average intensity projections of the Z-stack. To subtract background, the mode of signal measured outside of cells in each image was subtracted from the mean signal obtained from each cell.

### Spot fluorescence quantification in live-cell microscopy

The intensity of RNA spots was measured on a maximum intensity projection of the Z-stack. A region of interest was placed manually on a spot and another region of interest in the cytoplasm of the same cell. The maximum intensity measured for the spot was divided by the maximum intensity measured in the cytoplasm.

### Spot counting in live-cell microscopy

To count RNA spots in live-cell imaging, spots were identified in Z-stacks using U-FISH (Xu et al., 2025). To accommodate for the size of point-spread functions and movement of RNAs, detected spots were visually inspected and corrected within Napari. False-positive spots were manually deleted and RNA spots which moved between Z-slices to cause multiple detections were quantified as a single detected RNA. Remaining spots in close proximity (< 4 pixels in 3D distance) were averaged and interpreted as a single spot during additional data processing. To determine the number of spots per cell, cells were manually segmented in FIJI/ImageJ (Schindelin et al., 2012) and spots were assigned to cells based on their X/Y position.

### Single-molecule RNA fluorescence in situ hybridization (smRNA FISH)

Cultures were grown to a density of 0.8 to 1.5 × 107 cells/mL in Edinburgh Minimal Medium (EMM, MP Biochemicals LLC, 114110012). 2 × 108 cells were fixed by adding 16 % paraformaldehyde in PBS directly to the cell suspension to a final concentration of 2 % paraformaldehyde. Incubation was continued with shaking for 15 min at 30 °C and 15 min at room temperature. The fixation was quenched by pelleting the cells and resuspending them in 1 mL 50 mM ammonium chloride in PBS, incubating for 10 min, followed by three washes with PBS. If needed, cells were stored at this step at 4 °C. To remove the cell wall, cells were pelleted and resuspended in 1 mL of spheroplast buffer (0.1 M potassium phosphate, 20 mM vanadyl ribonuclease complex (NEB, S1402S), 20 μM beta-mercaptoethanol in PBS), to which 4 μL of 100T zymolyase (10 mg/mL; US Biological, Z1005) was added. Cells were kept at 30 °C until the cell walls were sufficiently digested, which was observed by testing small samples for osmotic lysis in water (∼70% lysed). Cells were washed three times with PBS to remove zymolyase, then incubated in 1 mL of 0.01 % Triton X-100 in PBS for 20 min to permeabilize the plasma membrane and washed three times with PBS. Priming for hybridization was performed by two washes with 10 % formamide in 2x saline sodium citrate (SSC) buffer (Thermo Fisher Scientific, AM9770). Each sample was split into two technical replicates, each containing 1× 108 cells. For each technical replicate, 25 ng of Stellaris RNA FISH probes (CAL Fluor red 610 probes targeting EGFP or Quasar 570 probes targeting sfGFPcp) and/or 50 ng of Stellaris RNA FISH probes (Quasar 570 probes targeting 24x MS2V6 (Tutucci et al., 2017);

Biosearch Technologies, LGC; kindly gifted from Robert Singer) were used. Probes were diluted in buffer F (20 % formamide, 10 mM sodium phosphate buffer (pH 7.2)) to a final volume of 50 μL, heated to 95 °C for 3 min and allowed to cool to room temperature before 50 µL 4x SSC buffer was added. Cell pellets were resuspended in this 100 μL hybridization solution and incubated overnight at 37 °C in the dark.

Following overnight incubation, cells were washed twice for 30 min each with 200 µL 10 % formamide in 2× SSC at 37 °C, followed by a 6 min incubation with 200 µL 2× SSC at room temperature, and lastly 10 min with 10 µL 4′,6-diamidino-2-phenylindole (DAPI; 1 µg/mL) in PBS at room temperature. Cells were washed once with PBS, resuspended in PBS, and stored in the dark at 4 °C until imaging. Cells were mounted in SlowFade Diamond Antifade Mountant (Thermo Fisher Scientific, S36972) using diethyl pyrocarbonate(DEPC)-treated slides and #1.5 glass coverslips.

### Fixed-cell microscopy

Cells were imaged with a Zeiss AxioImager M1 equipped with Xcite Fire LED illumination (Excelitas), a Zeiss Plan-APO 100x/1.45 oil objective and an ORCA-Flash4.0LT sCMOS camera (Hamamatsu). CAL610 and Quasar 570 FISH probes were imaged using a Chroma 49306 (ET–Red#1 FISH) and a Chroma 49304 (ET–Gold FISH) filter set, respectively. Cyan fluorescent protein (CFP) and DAPI were imaged using a Chroma 49001 and Chroma 49000 filter set, respectively. The images for each channel consisted of a 6 µm Z-stack containing 31 images at 0.2 µm intervals.

### smRNA FISH image analysis

Images were dark noise-subtracted and flatfielded using FIJI macros (Schindelin et al., 2012). Cell areas were segmented based on autofluorescence using Trainable WEKA segmentation (Arganda-Carreras et al., 2017) as a FIJI plugin, and RNA spots were detected using U-FISH as a napari plugin (Chiu et al., 2022; Xu et al., 2025). Following segmentation and spot detection, quality control checks and manual correction were performed in napari. Accuracy of spot detection was verified by manual counting spots in a subset of cells. In manual correction, cells close to debris, which would interfere with accurate spot quantification, and poorly segmented, overlapping, or non-intact cells were eliminated from the analysis along with all their respective RNAs.

To reduce the number of false-positive EGFP spots (Fig. S6B) for the co-localization analysis, an intensity filter was applied to the detected spots. The cutoff was determined by comparing the intensities of detected spots in a strain lacking EGFP mRNAs to the intensities of detected spots in a strain expressing EGFP mRNA. The chosen cutoff of 350 removed the majority of low-intensity false-positive spots while retaining the majority of spots detected in the strain with EGFP expression. For the co-localization analysis, the distance of the nearest neighbour of the opposite probe type was determined in 2D. Mutual nearest neighbour pairs were confirmed by reciprocal identification. Nearest neighbour pairs were designated as being overlapping if their inter-spot distance was below 300 nm.

## Acknowledgments

We thank Weihan Li, Dong Woo Hwang, and Robert Singer for advice on MS2-MCP imaging, the Singer lab for MS2V6 probes, Evelina Tutucci, Robert Singer, Atsushi Miyawaki, Aleksandar Vještica, Sophie Martin, Yoshinori Watanabe, and Jürg Bähler for plasmids, as well as Max Huynh and Himadri Sunam for help with creating vectors and strains.

## Data and resource availability

The likely most useful vectors have been deposited at Addgene (ID 255008-255017) and RIKEN BRC, as indicated in Table S2. All other vectors and yeast strains are available from the corresponding author upon request.

## Author contributions

Conceptualization, D.E.W. and S.H.; Investigation, S.C.T., S.G.C., and S.H.; Writing – Original Draft, S.H.; Writing – Review & Editing, D.E.W., S.C.T., S.G.C.; Visualization, S.G.C., S.H.; Supervision, S.H.; Funding Acquisition, S.H.

