## Supplementary material for "Establishing MS2-MCP-based single-molecule RNA visualization in *Schizosaccharomyces pombe*": Figures S1-S7, Tables S1, S2

#### Figure S1

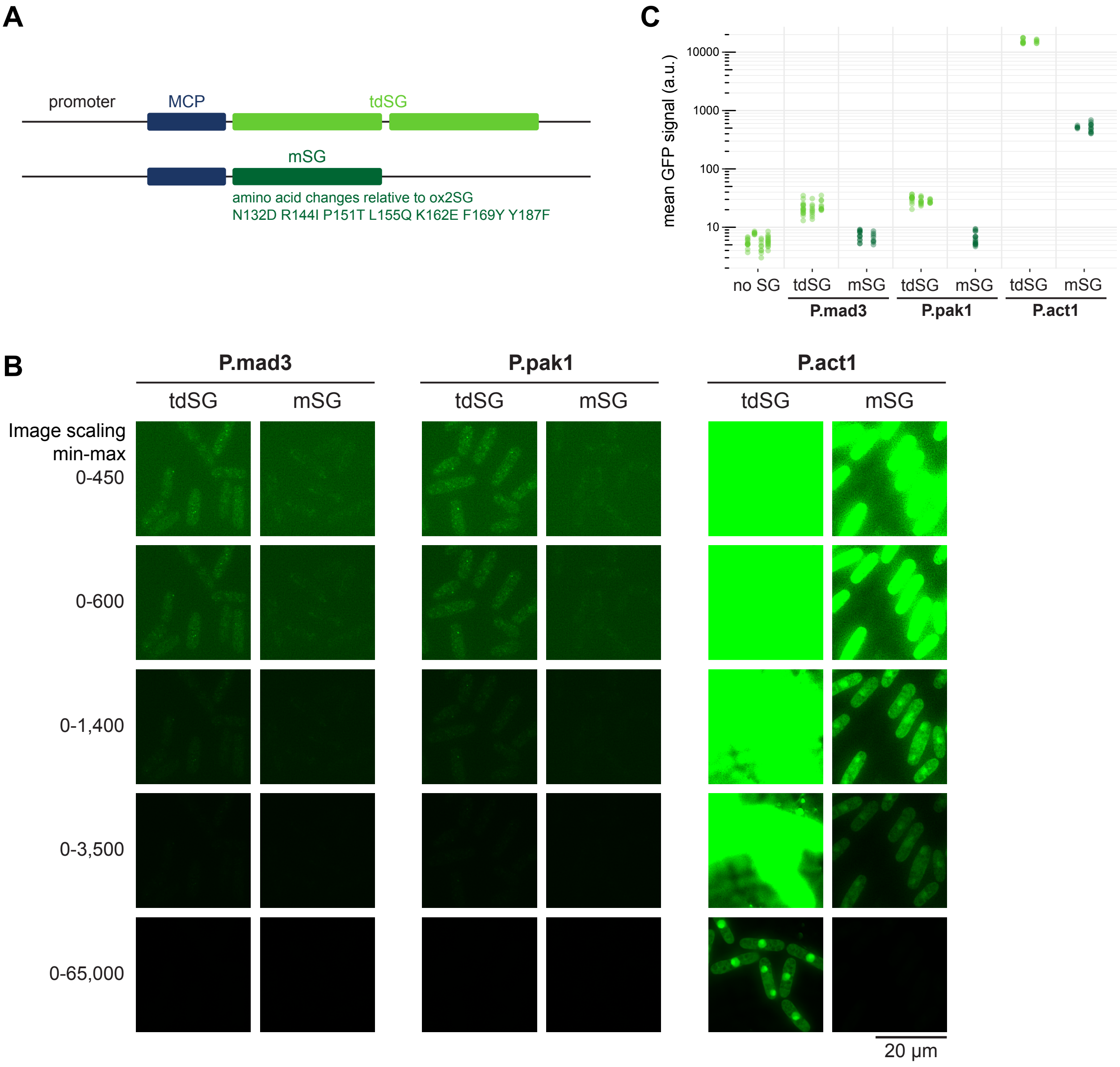


**Figure S1. Comparison between tandem and monomeric StayGold fused to MCP.**
**(A)** Schematic showing MCP-NLS-td8ox2SG (tandem SG, tdSG) and MCP-NLS-mSG (monomeric SG, mSG); mSG carries 7 amino acid changes relative to ox2SG. **(B)** Example images from strains expressing *mad2*-24xMS2 and MCP-NLS-tdSG or MCP-NLS-mSG from the *mad3*, *pak1*, or *act1* promoter. Image acquisition conditions were the same; each field of view is shown using five different scaling settings in order to capture the breadth of signal intensities. Example images for tdSG are the same as in Fig. 1. **(C)** The mean StayGold signal intensity in single cells was quantified; dots are individual cells, columns are technical or biological replicates. Note that data are displayed on a logarithmic scale. Signal intensity obtained from mSG is considerably less than half of that obtained from tdSG.

#### Figure S2

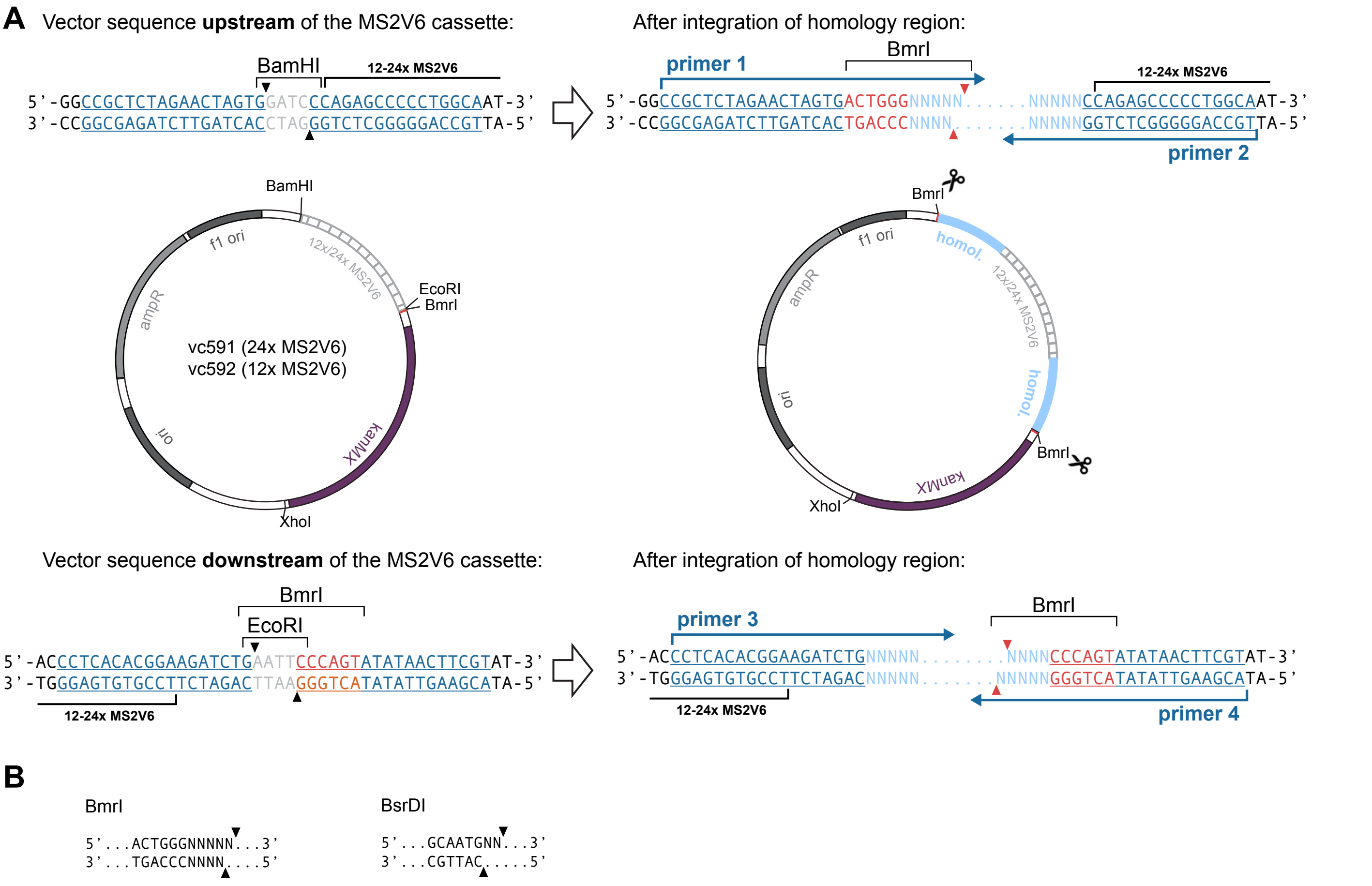


**Figure S2. Attaching homology regions to MS2V6 repeats.**
**(A)** Strategy for the integration of upstream and downstream homology regions (cyan), when BmrI is used as enzyme to cut out the piece to be transformed. The vectors already contain a BmrI site that overlaps with the EcoRI site downstream of the MS2V6 repeats. The upstream homology region is amplified with primers 1 and 2, the downstream homology region with primers 3 and 4. The PCR fragments are integrated into the BamHI/EcoRI-digested vector by Gibson assembly. Sequence regions: dark blue and underlined, regions of overlap with the vector for Gibson assembly; gray, nucleotides that become removed during Gibson assembly; orange, BmrI recognition site; cyan, inserted homology region; arrowheads: cut sites. In situations where a BmrI site is present in one of the homology regions, BsrDI sites can be introduced instead. **(B)** Type IIS restriction enzymes, such as BmrI and BsrDI, allow cleaving out the homology–MS2V6–homology fragment without leaving any traces of exogenous sequence.

### Figure S3

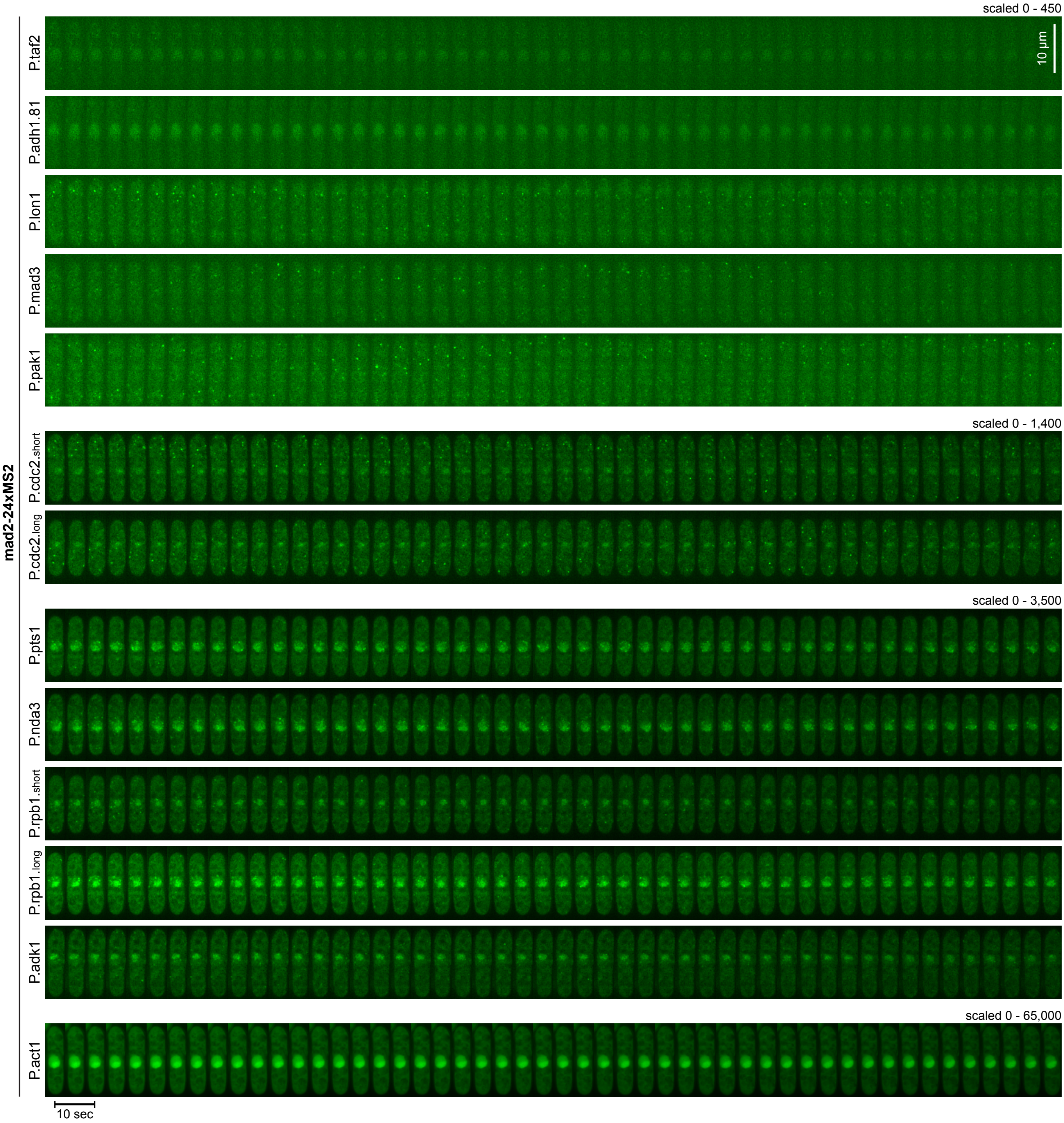


**Figure S3. Overview of all promoter-MCP-tdSG combinations tested for *mad2*-24xMS2 mRNA imaging.** Kymographs from live-cell imaging of the indicated strains expressing *mad2*-24xMS2. MCP-tdSG was expressed from the indicated promoters. Images are maximum intensity projections of the Z-stack. Note the different scaling settings for displaying the images, necessitated by the different expression levels.

#### Figure S4

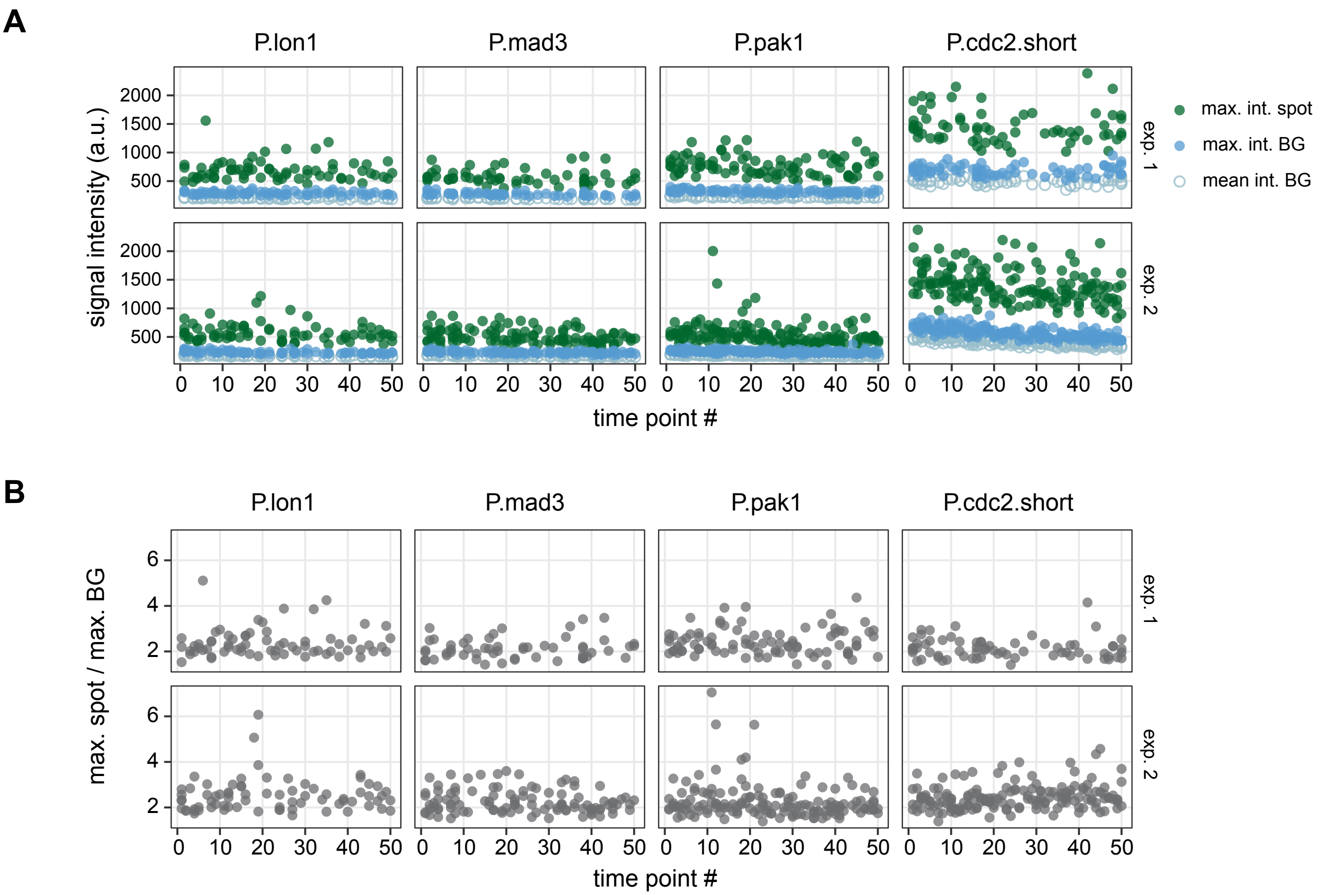


**Figure S4. Quantification of cytoplasmic RNA signals.**
Quantification of cytoplasmic dot-like signals (spots) and cytoplasmic background (BG) over time from two different experiments (exp. 1, exp. 2). Time points are spaced by 5 sec. **(A)** The maximum signal intensity is plotted for spots (green) and cytoplasmic background (blue); the mean intensity for the cytoplasmic background is shown as cyan circles. **(B)** Ratio between the maximum intensity of a spot and the maximum intensity of the cytoplasmic background in the same cell. The number of quantified spots per experiment and strain ranged from 56 to 90 for exp. 1 and 74 to 162 for exp. 2.

### Figure S5

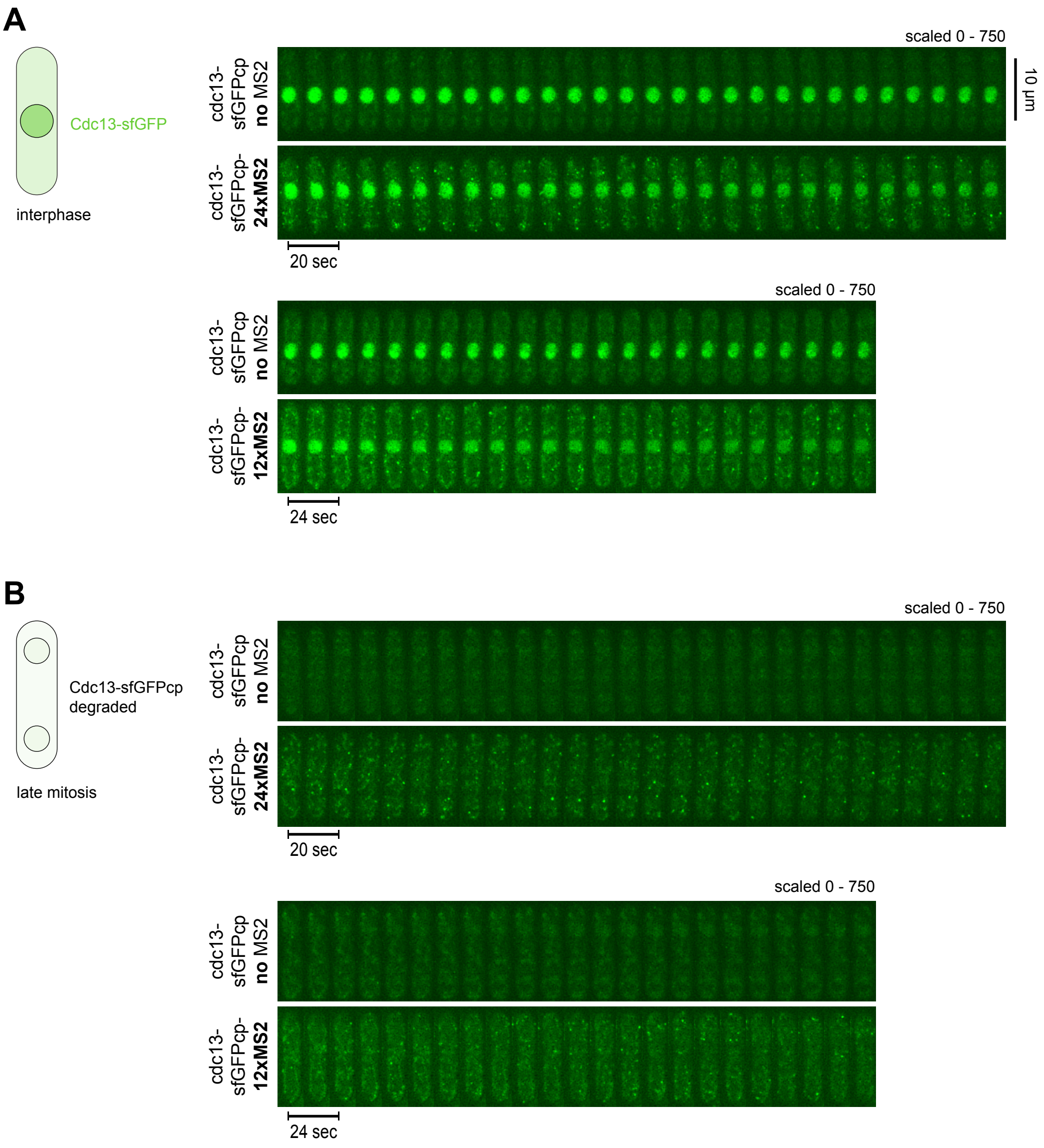


**Figure S5. Additional representative kymographs from cells expressing *cdc13*-sfGFPcp-MS2 and P.mad3-MCP-tdSG.** Kymographs from live-cell imaging of the indicated strains. A strain without integration of MS2 repeats is shown as control. Images were recorded every 5 sec for *cdc13*-sfGFPcp-24xMS2 and every 6 sec for *cdc13*-sfGFPcp-12xMS2; only every second image (10 sec and 12 sec, respectively) is shown. Images are maximum intensity projections of the Z-stack. **(A)** Cells in interphase with nuclear Cdc13-sfGFPcp signal. **(B)** Cells after nuclear division when the Cdc13-sfGFPcp protein has been degraded.

#### Figure S6

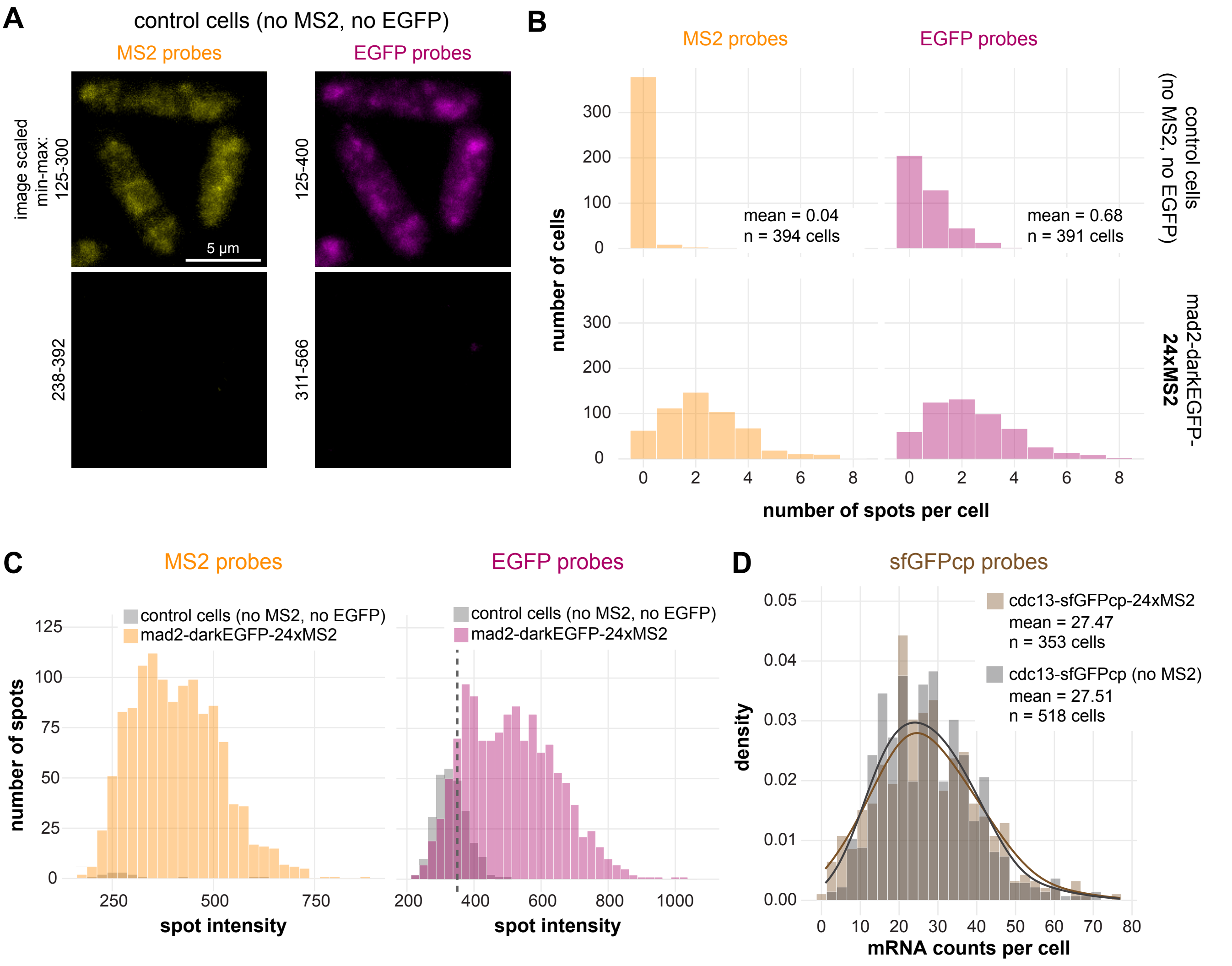


**Figure S6. Controls for RNA FISH.**
**(A)** Representative images (maximum intensity projections) of a control strain lacking MS2 and EGFP tags probed for both of these targets. Top: images scaled to make background visible; bottom: images scaled the same as shown in Fig. 4B. **(B)** Spot counts per cell for the control strain shown in (A). The bottom panels show the same data as in Fig. 4D. **(C)** Intensity of the spots detected in the control strain or in the *mad2*-darkEGFP-24xMS2 strain using MS2 or EGFP probes. The false positive signals in the control strain had weaker signal intensities. False positive signals were more common for the EGFP probes. To largely exclude these false positives, only EGFP spots with an intensity at the dashed line or higher were included in the co-localization analysis. **(D)** Comparison of mRNA counts per cell in samples probed for sfGFPcp for a strain expressing *cdc13*-S177S-sfGFPcp-24xMS2 and a strain expressing *cdc13*-S177S-sfGFPcp (no MS2). The data for the cdc13-sfGFPcp-24xMS2 strain is the same as in Fig. 4G.

#### Figure S7

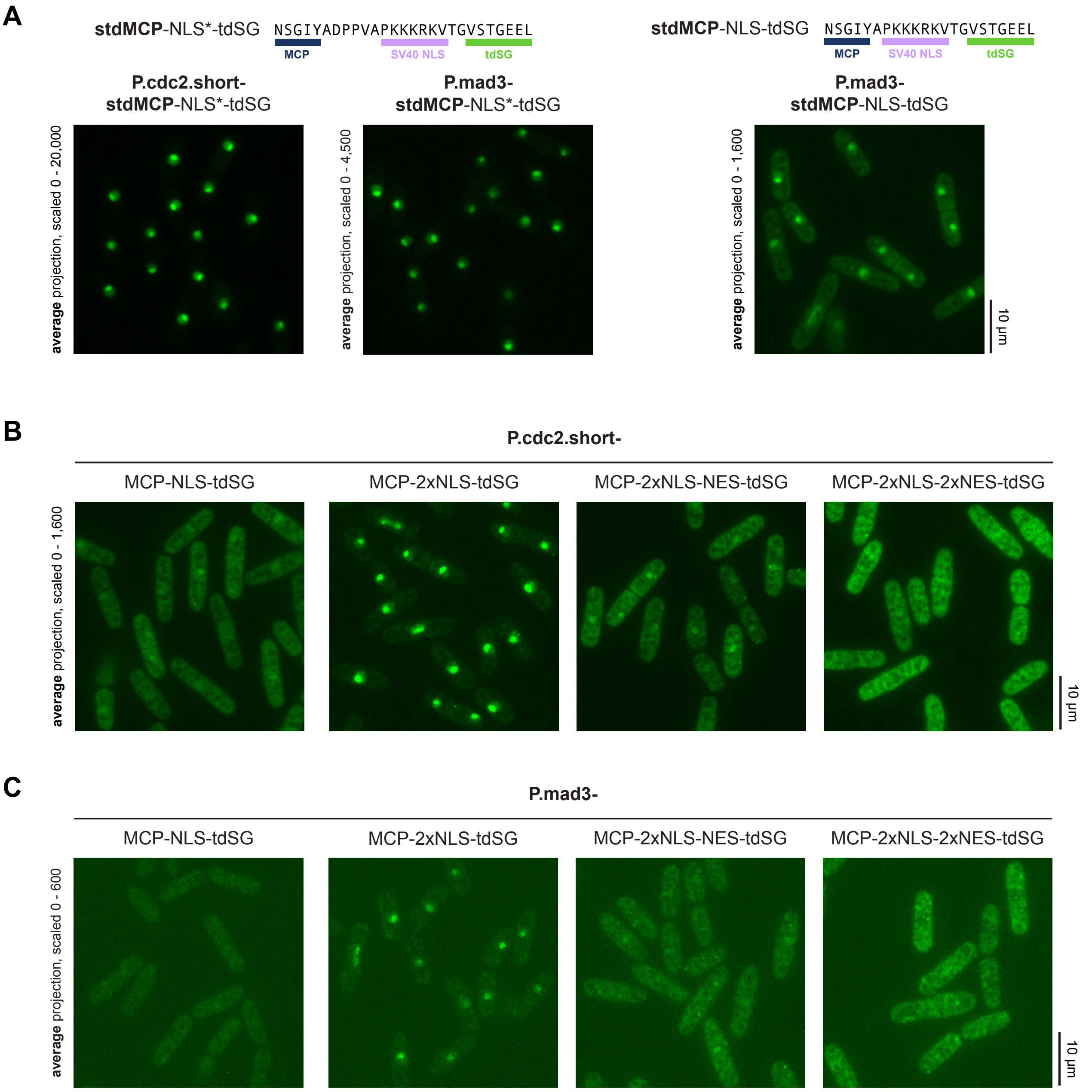


**Figure S7. Additional representative images for strains with different NLS/NES combinations.**
Representative overview images of the indicated strains expressing *mad2*-24xMS2 and MCP-tdSG with different number of NLSs or NLS/NES combinations. Shown are average intensity projections (not maximum intensity projections) to adequately represent the nucleo-cytoplasmic ratios. Note the different scaling settings for displaying the images, necessitated by the different expression levels and nuclear enrichment. **(A)** Cells expressing synonymized tandem MCP (stdMCP) with either a longer (NLS*) or a shorter (NLS) linker between MCP and NLS. **(B)** Cells expressing MCP-tdSG from the short version of the *cdc2* promoter. **(C)** Cells expressing MCP-tdSG from the *mad3* promoter.

#### Supplemental Movies

**Movie S1. Live-cell imaging of *mad2*-24xMS2 with P.lon1-MCP-tdSG**

Cells expressing MCP-tdSG from the *lon1* promoter. **Top**: cells expressing *mad2*-24xMS2; **bottom**: cells without MS2 stem-loops as control. Z-stacks were acquired every 5 sec; a maximum intensity projection is shown.

**Movie S2. Live-cell imaging of *mad2*-24xMS2 with P.mad3-MCP-tdSG**

Cells expressing MCP-tdSG from the *mad3* promoter. **Top**: cells expressing *mad2*-24xMS2; **bottom**: cells without MS2 stem-loops as control. Z-stacks were acquired every 5 sec; a maximum intensity projection is shown.

**Movie S3. Live-cell imaging of *mad2*-24xMS2 with P.pak1-MCP-tdSG**

Cells expressing MCP-tdSG from the *pak1 (shk1)* promoter. **Top**: cells expressing *mad2*-24xMS2; **bottom**: cells without MS2 stem-loops as control. Z-stacks were acquired every 5 sec; a maximum intensity projection is shown.

**Movie S4. Live-cell imaging of *mad2*-24xMS2 with P.cdc2.short-MCP-tdSG**

Cells expressing MCP-tdSG from the short *cdc2* promoter. **Top**: cells expressing *mad2*-24xMS2; **bottom**: cells without MS2 stem-loops as control. Z-stacks were acquired every 5 sec; a maximum intensity projection is shown.

**Movie S5. Live-cell imaging of *cdc13*-sfGFPcp-24xMS2 with P.mad3-MCP-tdSG**

Cells expressing MCP-tdSG from the *mad3* promoter. **Top**: cells expressing *cdc13*-sfGFPcp-24xMS2; bottom: cells expressing *cdc13*-sfGFPcp (no MS2) as control. Z-stacks were acquired every 5 sec; a maximum intensity projection is shown. Note that both MCP-tdSG and Cdc13-sfGFPcp are visible in the green channel.

**Movie S6. Live-cell imaging of *cdc13*-sfGFPcp-12xMS2 with P.mad3-MCP-tdSG**

Cells expressing MCP-tdSG from the *mad3* promoter. **Top**: cells expressing *cdc13*-sfGFPcp-12xMS2; **bottom**: cells expressing *cdc13*-sfGFPcp (no MS2) as control. Z-stacks were acquired every 6 sec; a maximum intensity projection is shown. Note that both MCP-tdSG and Cdc13-sfGFPcp are visible in the green channel.

**Movie S7. Live-cell imaging of *rad21*-24xMS2 with P.mad3-MCP-tdSG**

Cells expressing MCP-tdSG from the *mad3* promoter as well as *rad21*-24xMS2. Z-stacks were acquired every 5 min; a maximum intensity projection is shown. The *rad21* (cohesin) mRNA shows cell cycle-dependent expression just prior to septation and cell division.

**Table S1.** ***S. pombe*** **strains.**

| Name | Mating type | Genotype | Figure(s) | Simplified genotype |
|---|---|---|---|---|
| SX293 | *h-* | *leu1- ura4+::P.taf2(244-1)-MCP-NLS-td8ox2StayGold-Scer\T.ADH1 mad2-ntG606A-ymEGFP-Y66L-3UTR.mad2-G0G-24xMS2V6* | 1, S3 | *P.taf2-MCP-NLS-tdSG mad2-24xMS2* |
| SX291 | *h-* | *leu1- ura4+::P.lon1(451-1)-MCP-NLS-td8ox2StayGold-Scer\T.ADH1 mad2-ntG606A-ymEGFP-Y66L-3UTR.mad2-G0G-24xMS2V6* | 1, 3, S3, S4 | *P.lon1-MCP-NLS-tdSG mad2-24xMS2* |
| SX258 | *?* | *leu1- ura4+::P.mad3(717-1)-MCP-NLS-td8ox2StayGold-Scer\T.ADH1 mad2-ntG606A-ymEGFP-Y66L-3UTR.mad2-G0G-24xMS2V6* | 1, 3, 4, S1, S3, S4, S6, S7 | *P.mad3-MCP-NLS-tdSG mad2-24xMS2* |
| SX716 | *h-* | *leu1- ura4+::P.pak1-MCP-NLS-td8ox2StayGold-Scer\T.ADH1 mad2-ntG606A-ymEGFP-Y66L-3UTR.mad2-G0G-24xMS2V6* | 1, 3, S1, S3, S4 | *P.pak1-MCP-NLS-tdSG mad2-24xMS2* |
| SX717 | *h-* | *leu1- ura4+::P.adh1.81-MCP-NLS-td8ox2StayGold-Scer\T.ADH1 mad2-ntG606A-ymEGFP-Y66L-3UTR.mad2-G0G-24xMS2V6* | 1, S3 | *P.adh1.81-MCP-NLS-tdSG mad2-24xMS2* |
| SX296 | *h-* | *leu1- ura4+::P.cdc2(355-1)-MCP-NLS-td8ox2StayGold-Scer\T.ADH1 mad2-ntG606A-ymEGFP-Y66L-3UTR.mad2-G0G-24xMS2V6* | 1, 3, 6, S3, S4, S7 | *P.cdc2.short-MCP-NLS-tdSG mad2-24xMS2* |
| SX297 | *h-* | *leu1- ura4+::P.cdc2(913-1)-MCP-NLS-td8ox2StayGold-Scer\T.ADH1 mad2-ntG606A-ymEGFP-Y66L-3UTR.mad2-G0G-24xMS2V6* | 1, S3 | *P.cdc2.long-MCP-NLS-tdSG mad2-24xMS2* |
| SX292 | *h-* | *leu1- ura4+::P.pts1(378-1)-MCP-NLS-td8ox2StayGold-Scer\T.ADH1 mad2-ntG606A-ymEGFP-Y66L-3UTR.mad2-G0G-24xMS2V6* | 1, S3 | *P.pts1-MCP-NLS-tdSG mad2-24xMS2* |
| SX283 | *h-* | *leu1- ura4+::P.nda3(620-1)-MCP-NLS-td8ox2StayGold-Scer\T.ADH1 mad2-ntG606A-ymEGFP-Y66L-3UTR.mad2-G0G-24xMS2V6* | 1, S3 | *P.nda3-MCP-NLS-tdSG mad2-24xMS2* |
| SX284 | *h-* | *leu1- ura4+::P.rpb1(214-1)-MCP-NLS-td8ox2StayGold-Scer\T.ADH1 mad2-ntG606A-ymEGFP-Y66L-3UTR.mad2-G0G-24xMS2V6* | 1, S3 | *P.rpb1.short-MCP-NLS-tdSG mad2-24xMS2* |
| SX285 | *h-* | *leu1- ura4+::P.rpb1(647-1)-MCP-NLS-td8ox2StayGold-Scer\T.ADH1 mad2-ntG606A-ymEGFP-Y66L-3UTR.mad2-G0G-24xMS2V6* | 1, S3 | *P.rpb1.long-MCP-NLS-tdSG mad2-24xMS2* |
| SX282 | *h-* | *leu1- ura4+::P.adk1(338-1)-MCP-NLS-td8ox2StayGold-Scer\T.ADH1 mad2-ntG606A-ymEGFP-Y66L-3UTR.mad2-G0G-24xMS2V6* | 1, S3 | *P.adk1-MCP-NLS-tdSG mad2-24xMS2* |
| SX250 | *?* | *leu1- ura4+::P.act1(822-1)-MCP-NLS-td8ox2StayGold-Scer\T.ADH1 mad2-ntG606A-ymEGFP-Y66L-3UTR.mad2-G0G-24xMS2V6* | 1, S1, S3 | *P.act1-MCP-NLS-tdSG mad2-24xMS2* |
| SX257 | *?* | *leu1- ura4+::P.mad3(717-1)-MCP-NLS-mStayGold-Scer\T.ADH1 mad2-ntG606A-ymEGFP-Y66L-3UTR.mad2-G0G-24xMS2V6* | S1 | *P.mad3-MCP-NLS-mSG mad2-24xMS2* |
| SX259 | *?* | *leu1- ura4+::P.pak1(630-1)-MCP-NLS-mStayGold-Scer\T.ADH1 mad2-ntG606A-ymEGFP-Y66L-3UTR.mad2-G0G-24xMS2V6* | S1 | *P.pak1-MCP-NLS-mSG mad2-24xMS2* |
| SX249 | *?* | *leu1- ura4+::P.act1(822-1)-MCP-mStayGold-Scer\T.ADH1 mad2-ntG606A-ymEGFP-Y66L-3UTR.mad2-G0G-24xMS2V6* | S1 | *P.act1-MCP-NLS-mSG mad2-24xMS2* |
| SX279 | *?* | *leu1- ura4-D18 mad2-ntG606A-ymEGFP-Y66L-3UTR.mad2-G0G-24xMS2V6* | 3, S1 | *mad2-24xMS2* |
| SX280 | *?* | *leu1- ura4-D18 mad2-ntG606A-ymEGFP-Y66L-3UTR.mad2-G0G-12xMS2V6* | S1 | *mad2-12xMS2* |
| SX724 | *h-* | *leu1- ura4+::P.lon1-MCP-NLS-td8ox2StayGold-Scer\T.ADH1* | 3 | *P.lon1-MCP-NLS-tdSG* |
| SX253 | *h-* | *leu1- ura4+::P.mad3(717-1)-MCP-NLS-td8ox2StayGold-Scer\T.ADH1* | 3, S6 | *P.mad3-MCP-NLS-tdSG* |
| SX725 | *h-* | *leu1- ura4+::P.pak1-MCP-NLS-td8ox2StayGold-Scer\T.ADH1* | 3 | *P.pak1-MCP-NLS-tdSG* |

| Name | Mating type | Genotype | Figure(s) | Simplified genotype |
|---|---|---|---|---|
| SX723 | *h-* | *leu1- ura4+::P.cdc2(355-1)-MCP-NLS-td8ox2StayGold-Scer\T.ADH1* | 3 | *P.cdc2.short-MCP-NLS-tdSG* |
| SX761 | *h-* | *leu1-32::P.cdc13-cdc13-S177S-sfGFPcp:leu1+ ura4+::P.mad3(717-1)-MCP-NLS-td8ox2StayGold-Scer\T.ADH1* | 3, S5, S6 | *P.mad3-MCP-NLS-mSG cdc13-sfGFPcp* (exogenous) |
| SX289 | *h-* | *leu1-32::P.cdc13-cdc13-S177S-sfGFPcp-3UTR.cdc13-C332C-24xMS2V6:leu1+ ura4+::P.mad3(717-1)-MCP-NLS-td8ox2StayGold-Scer\T.ADH1* | 3, 4, S5, S6 | *P.mad3-MCP-NLS-mSG cdc13-sfGFPcp-24xMS2(332)* (exogenous) |
| SX287 | *h-* | *leu1-32::P.cdc13-cdc13-S177S-sfGFPcp-3UTR.cdc13-C332C-12xMS2V6:leu1+ ura4+::P.mad3(717-1)-MCP-NLS-td8ox2StayGold-Scer\T.ADH1* | 3, S5 | *P.mad3-MCP-NLS-mSG cdc13-sfGFPcp-12xMS2(332)* (exogenous) |
| SX286 | *h-* | *leu1-32::P.cdc13-cdc13-S177S-sfGFPcp-3UTR.cdc13-A205A-12xMS2V6:leu1+ ura4+::P.mad3(717-1)-MCP-NLS-td8ox2StayGold-Scer\T.ADH1* | 3, S5 | *P.mad3-MCP-NLS-mSG cdc13-sfGFPcp-12xMS2(205)* (exogenous) |
| SX288 | *h-* | *leu1-32::P.cdc13-cdc13-S177S-sfGFPcp-3UTR.cdc13-A205A-24xMS2V6:leu1+ ura4+::P.mad3(717-1)-MCP-NLS-td8ox2StayGold-Scer\T.ADH1* | S5 | *P.mad3-MCP-NLS-mSG cdc13-sfGFPcp-24xMS2(205)* (exogenous) |
| SX795 | *h-* | *leu1-32 ade6+::P.rad21(1452-1)-rad21-3UTR.rad21-A70A24xMS2V6 ura4+::P.mad3(717-1)-MCP-NLS-td8ox2StayGold-Scer\T.ADH1* | 5 | *P.mad3-MCP-NLS-tdSG rad21-24xMS2* (exogenous) |
| SX720 | *h-* | *leu1- ura4+::P.cdc2(355-1)-MCP-2xNLS-td8ox2StayGold-Scer\T.ADH1 mad2-ntG606A-ymEGFP-Y66L-3UTR.mad2-G0G-24xMS2V6* | 6, S7 | *P.cdc2.short-MCP-2xNLS-tdSG mad2-24xMS2* |
| SX721 | *h-* | *leu1- ura4+::P.cdc2(355-1)-MCP-2xNLS-NES-td8ox2StayGold-Scer\T.ADH1 mad2-ntG606A-ymEGFP-Y66L-3UTR.mad2-G0G-24xMS2V6* | 6, S7 | *P.cdc2.short-MCP-2xNLS-NES-tdSG mad2-24xMS2* |
| SX722 | *h-* | *leu1- ura4+::P.cdc2(355-1)-MCP-2xNLS-2xNES-td8ox2StayGold-Scer\T.ADH1 mad2-ntG606A-ymEGFP-Y66L-3UTR.mad2-G0G-24xMS2V6* | 6, S7 | *P.cdc2.short-MCP-2xNLS-2xNES-tdSG mad2-24xMS2* |
| SX719 | *h-* | *leu1- ura4+::P.cdc2(355-1)-stdMCP-NLS-td8ox2StayGold-Scer\T.ADH1 mad2-ntG606A-ymEGFP-Y66L-3UTR.mad2-G0G-24xMS2V6* | 6, S7 | *P.cdc2.short-stdMCP-NLS*-tdSG mad2-24xMS2* |
| SX712 | *h-* | *leu1- ura4+::P.mad3(717-1)-MCP-2xNLS-td8ox2StayGold-Scer\T.ADH1 mad2-ntG606A-ymEGFP-Y66L-3UTR.mad2-G0G-24xMS2V6* | S7 | *P.mad3-MCP-2xNLS-tdSG mad2-24xMS2* |
| SX733 | *h-* | *leu1- ura4+::P.mad3-MCP-2xNLS-NES-td8ox2StayGold-Scer\T.ADH1 mad2-ntG606A-ymEGFP-Y66L-3UTR.mad2-G0G-24xMS2V6* | S7 | *P.mad3-MCP-2xNLS-NES-tdSG mad2-24xMS2* |
| SX714 | *h-* | *leu1- ura4+::P.mad3(717-1)-MCP-2xNLS-2xNES-td8ox2StayGold-Scer\T.ADH1 mad2-ntG606A-ymEGFP-Y66L-3UTR.mad2-G0G-24xMS2V6* | S7 | *P.mad3-MCP-2xNLS-2xNES-tdSG mad2-24xMS2* |
| SX718 | *h-* | *leu1- ura4+::P.mad3(717-1)-stdMCP-NLS-td8ox2StayGold-Scer\T.ADH1 mad2-ntG606A-ymEGFP-Y66L-3UTR.mad2-G0G-24xMS2V6* | S7 | *P.mad3-stdMCP-NLS*-tdSG mad2-24xMS2* |
| SX767 | *?* | *leu1- mad2-ntG606A-ymEGFP-Y66L-3UTR.mad2-G0G-24xMS2V6 ura4+::P.mad3(717-1)-stdMCP-NLS-td8ox2StayGold-Scer\T.ADH1* | S7 | *P.mad3-stdMCP-NLS-tdSG mad2-24xMS2* |

#### Table S2. Vectors.

pUra4 vectors are derived from pUra4AfeI, Addgene 133467, doi: 10.1242/jcs.240754
The pAde6 vector is derived from pAde6PmeI, Addgene 133468, doi: 10.1242/jcs.240754
MCP (MS2 coat protein) is from Addgene 104394, doi: 10.1038/nmeth.4502.
stdMCP (synonymized tandem MCP) is from Addgene 98916, doi: 10.1101/gad.259358.115
Monomeric StayGold (mSG) is from RIKEN RDB20214, doi: 10.1038/s41592-023-02085-6
TandemStayGold (tdSG) is from RIKEN RDB20227, doi: 10.1038/s41592-023-02085-6
24xMS2V6 is from Addgene 104393, doi: 10.1038/nmeth.4502
12xMS2V6 is from Addgene 104392, doi: 10.1038/nmeth.4502
pDUAL vectors are based on Matsuyama, Yoshida *et al.*, doi: 10.1002/yea.1181

| ID | name | insert | notes | deposited |
|---|---|---|---|---|
| vc547 | pUra4-P.mad3-MCP-NLS-mSG | *P.mad3(717-1)-MCP-NLS-mStayGold-Scer\T.ADH1* | | N |
| vc548 | pUra4-P.mad3-MCP-NLS-tdSG | *P.mad3(717-1)-MCP-NLS-td8ox2StayGold-Scer\T.ADH1* | | Y |
| vc549 | pUra4-P.pak1-MCP-NLS-mSG | *P.pak1(630-1)-MCP-NLS-mStayGold-Scer\T.ADH1* | | N |
| vc582 | pUra4-P.act1-MCP-mSG | *P.act1(822-1)-MCP-GGGGS-mStayGold-Scer\T.ADH1* | | N |
| vc583 | pUra4-P.act1-MCP-NLS-tdSG | *P.act1(822-1)-MCP-NLS-td8ox2StayGold-Scer\T.ADH1* | | N |
| vc591 | pET264-pUC_24xMS2V6_homIns | *24xMS2V6* | | Y |
| vc592 | pET251-pUC_12xMS2V6_homIns | *12xMS2V6* | | Y |
| vc603 | pUra4-P.adk1-MCP-NLS-tdSG | *P.adk1(338-1)-MCP-NLS-td8ox2StayGold-Scer\T.ADH1* | | N |
| vc604 | pUra4-P.nda3-MCP-NLS-tdSG | *P.nda3(620-1)-MCP-NLS-td8ox2StayGold-Scer\T.ADH1* | | N |
| vc605 | pUra4-P.rpb1.short-MCP-NLS-tdSG | *P.rpb1(214-1)-MCP-NLS-td8ox2StayGold-Scer\T.ADH1* | | N |
| vc606 | pUra4-P.rpb1.long-MCP-NLS-tdSG | *P.rpb1(647-1)-MCP-NLS-td8ox2StayGold-Scer\T.ADH1* | | N |
| vc607 | pDUAL-cdc13-sfGFP-12xMS2V6_3p-205 | *P.cdc13-cdc13-sfGFPcp-12xMS2V6 (pos. 205 in 3' UTR)* | | N |
| vc608 | pDUAL-cdc13-sfGFP-12xMS2V6_3p-332 | *P.cdc13-cdc13-sfGFPcp-12xMS2V6 (pos. 332 in 3' UTR)* | | N |
| vc609 | pDUAL-cdc13-sfGFP-24xMS2V6_3p-305 | *P.cdc13-cdc13-sfGFPcp-24xMS2V6 (pos. 205 in 3' UTR)* | | N |
| vc610 | pDUAL-cdc13-sfGFP-24xMS2V6_3p-332 | *P.cdc13-cdc13-sfGFPcp-24xMS2V6 (pos. 332 in 3' UTR)* | | N |

| ID | name | insert | notes | depo-sited |
|---|---|---|---|---|
| vc617 | pUra4-P.cdc2.short-MCP-NLS-tdSG | *P.cdc2(355-1)-MCP-NLS-td8ox2StayGold-Scer\T.ADH1* | requires limited digest since AfeI site present in *cdc2* promoter | N |
| vc618 | pUra4-P.cdc2.long-MCP-NLS-tdSG | *P.cdc2(913-1)-MCP-NLS-td8ox2StayGold-Scer\T.ADH1* | requires limited digest since AfeI site present in *cdc2* promoter | N |
| vc619 | pUra4-P.lon1-MCP-NLS-tdSG | *P.lon1(451-1)-MCP-NLS-td8ox2StayGold-Scer\T.ADH1* | | Y |
| vc620 | pUra4-P.pts1-MCP-NLS-tdSG | *P.pts1(378-1)-MCP-NLS-td8ox2StayGold-Scer\T.ADH1* | | N |
| vc621 | pUra4-P.taf2-MCP-NLS-tdSG | *P.taf2(244-1)-MCP-NLS-td8ox2StayGold-Scer\T.ADH1* | | N |
| vc633 | pUra4-P.pak1-MCP-NLS-tdSG | *P.pak1(630-1)-MCP-NLS-td8ox2StayGold-Scer\T.ADH1* | *pak1* is an alternative gene name for *shk1* | Y |
| vc634 | pUra4-P.adh1.81-MCP-NLS-tdSG | *P.adh1.81-MCP-NLS-td8ox2StayGold-Scer\T.ADH1* | | N |
| vc640 | pUra4-P.mad3-MCP-2xNLS-tdSG | *P.mad3(717-1)-MCP-2xNLS-td8ox2StayGold-Scer\T.ADH1* | | N |
| vc641 | pUra4-P.mad3-MCP-2xNLS-NES-tdSG | *P.mad3(717-1)-MCP-2xNLS-NES-td8ox2StayGold-Scer\T.ADH1* | | Y |
| vc642 | pUra4-P.mad3-MCP-2xNLS-2xNES-tdSG | *P.mad3(717-1)-MCP-2xNLS-2xNES-td8ox2StayGold-Scer\T.ADH1* | | Y |
| vc643 | pUra4-P.cdc2.short-MCP-2xNLS-tdSG | *P.cdc2(355-1)-MCP-2xNLS-td8ox2StayGold-Scer\T.ADH1* | requires limited digest since AfeI site present in *cdc2* promoter | N |
| vc644 | pUra4-P.cdc2.short-MCP-2xNLS-NES-tdSG | *P.cdc2(355-1)-MCP-2xNLS-NES-td8ox2StayGold-Scer\T.ADH1* | requires limited digest since AfeI site present in *cdc2* promoter | N |
| vc645 | pUra4-P.cdc2.short-MCP-2xNLS-2xNES-tdSG | *P.cdc2(355-1)-MCP-2xNLS-2xNES-td8ox2StayGold-Scer\T.ADH1* | requires limited digest since AfeI site present in *cdc2* promoter | N |

| ID | name | insert | notes | depo-sited |
|---|---|---|---|---|
| vc646 | pUra4-P.mad3-stdMCP-NLS*-tdSG | *P.mad3(717-1)-stdMCP-NLS*-td8ox2StayGold-Scer\T.ADH1* | | N |
| vc647 | pUra4-P.cdc2.short-stdMCP-NLS*-tdSG | *P.cdc2(355-1)-stdMCP-NLS*-td8ox2StayGold-Scer\T.ADH1* | requires limited digest since AfeI site present in *cdc2* promoter | N |
| vc655 | pUra4-P.cdc2.short(AfeImut)-MCP-NLS-tdSG | *P.cdc2(355-1,AfeImut)-MCP-NLS-td8ox2StayGold-Scer\T.ADH1* | AfeI site within *cdc2* promoter mutated by removing one nucleotide | Y |
| vc669 | pUra4-P.mad3-stdMCP-NLS-tdSG | *P.mad3(717-1)-stdMCP-NLS-td8ox2StayGold-Scer\T.ADH1* | | Y |
| vc700 | pUra4-P.cdc2.short(AfeImut)-MCP-2xNLS-NES-tdSG | *P.cdc2(355-1,AfeImut)-MCP-2xNLS-NES-td8ox2StayGold-Scer\T.ADH1* | AfeI site within *cdc2* promoter mutated by removing one nucleotide | Y |
| vc701 | pUra4-P.cdc2.short(AfeImut)-MCP-2xNLS-2xNES-tdSG | *P.cdc2(355-1,AfeImut)-MCP-2xNLS-2xNES-td8ox2StayGold-Scer\T.ADH1* | AfeI site within *cdc2* promoter mutated by removing one nucleotide | Y |
| vc724 | pAde6-P.rad21.long-rad21-24xMS2V6_A70 | *P.rad21(1452-1)-rad21-3UTR.rad21-A70A24xMS2V6* | | N |
