## Supplementary figures and images for "Establishing MS2-MCP-based single-molecule RNA visualization in *Schizosaccharomyces pombe*"

### Movie S1

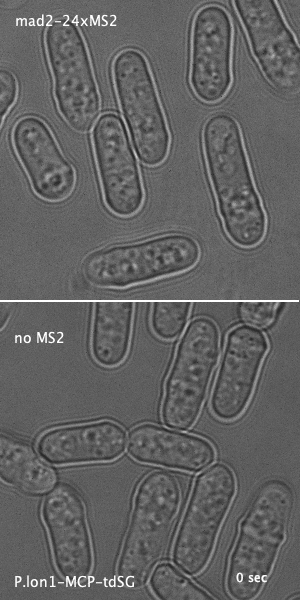

### Movie S2

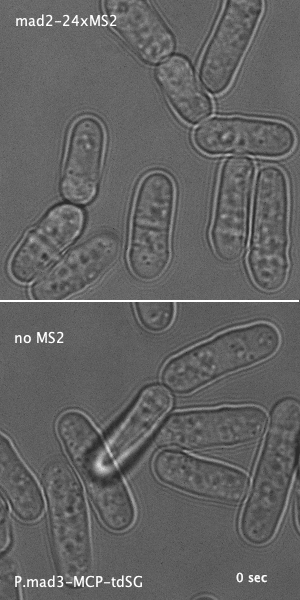

### Movie S3

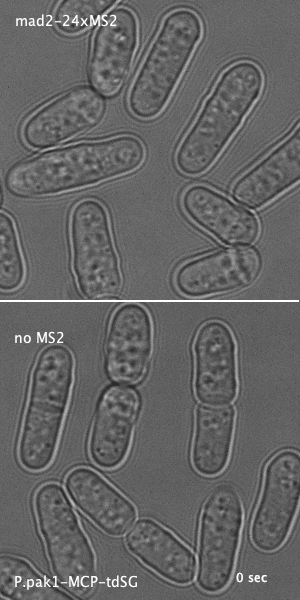

### Movie S4

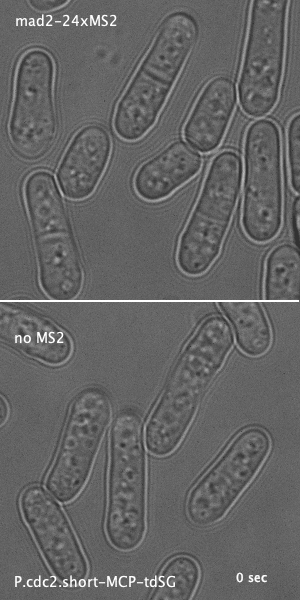

### Movie S5

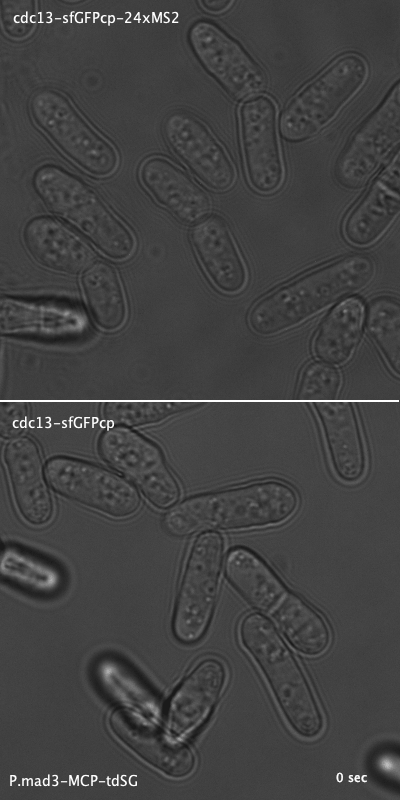

### Movie S6

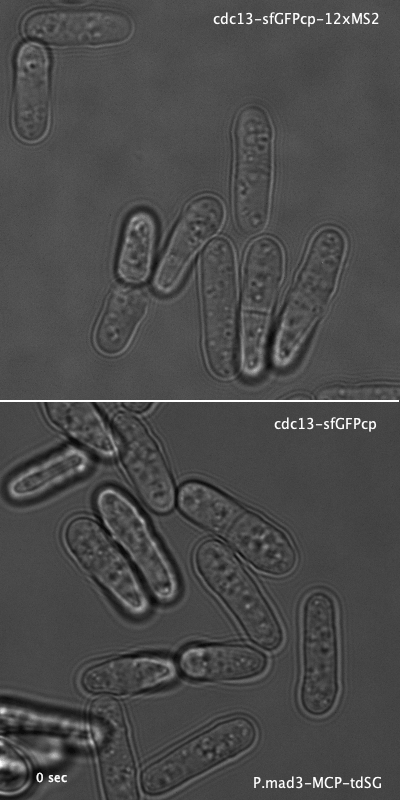

### Movie S7

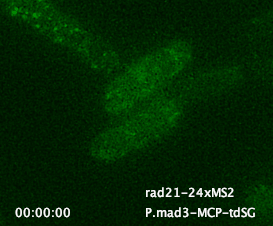
